# Dysregulated flow responses drive pathogenesis in a cellular model of Sturge–Weber syndrome

**DOI:** 10.64898/2026.06.02.729175

**Authors:** Kimihiko Banno, Ryuji Yokokawa, Jered Myslinski, Kazuaki Maruyama, Shota Watanabe, Junko Yoshida, Yoshikazu Kameda, Akiyoshi Kakita, Masaki Takao, Masaki Iwasaki, Akihiro Shindo, Atsuya Shirono, Yoshiaki Okada, Keiichiro Suzuki, Knut Woltjen, Takashi Hato, Yoshihiro Ujihara, Kyoji Horie

## Abstract

Vascular diseases comprise a broad spectrum of conditions associated with organ dysfunction, yet how endothelial responses to mechanical forces drive their pathogenesis remains unclear. Sturge–Weber syndrome (SWS), a flow-associated vascular malformation caused by somatic GNAQ mutations, provides a model to investigate this question. Here, using isogenic induced pluripotent stem cell–derived endothelial cells (ECs), perfusable vascular models, and simplified systems under defined flow conditions, we show that SWS ECs exhibited maladaptive responses to laminar flow, including impaired flow alignment, reduced CLDN5 expression, and decreased transendothelial electrical resistance. Importantly, three-dimensional acoustic microscopy uncovered disturbed surface changes in SWS ECs, potentially contributing to changes in mechanosensing signaling. Transcriptomic and chromatin accessibility analyses revealed dysregulated flow-responsive gene programs, some of which were validated in human SWS samples. These findings provide insights into flow-dependent endothelial dysfunction and a framework for therapeutic modulation in SWS and other vascular disorders.

## Introduction

Vascular diseases encompass a wide range of conditions that lead to dysfunction in various organs^1,2^. Despite their diversity, the contribution of endothelial responses to mechanical forces in disease pathophysiology remains insufficiently explored^3–5^. Vascular malformations, in particular, are classified into slow- and fast-flow lesions according to the ISSVA classification^6^; however, disease models that recapitulate pathological cellular processes, including alterations in cell shape, under physiologically relevant perfusion conditions remain scarce^7–9^.

Sturge–Weber syndrome (SWS) is a vascular malformation caused by a somatic activating mutation in *GNAQ* (c.548G>A, p.R183Q), leading to abnormal venous-like vasculature in the brain^10^. These vascular abnormalities are associated with severe clinical manifestations, including intractable epilepsy and progressive neurological impairment, highlighting the critical need to understand their underlying pathogenesis^11,12^. Although its genetic basis has been established, how this mutation gives rise to the characteristic vascular morphology remains unclear^13^. *GNAQ* encodes the Gαq subunit, which is classically known to mediate G protein–coupled receptor signaling^14^; however, recent evidence indicates that Gq/G11 signaling can also be activated by mechanical forces in endothelial cells (ECs)^15,16^, raising the possibility that altered responses to shear stress contribute to disease pathophysiology. We therefore hypothesized that modeling SWS under perfusion conditions would provide new insights into its underlying mechanisms.

In this study, we employed genome-edited induced pluripotent stem cells (iPSCs) modeling SWS to generate vascular networks from iPSC-derived ECs on-chip, which were then subjected to perfusion and analyzed using single-cell RNA-sequencing. Our findings reveal that responses to perfusion differ between SWS and wild-type cells. SWS ECs show altered morphological adaptation to pulsatile laminar flow, impaired endothelial barrier function, and distinct transcriptomic changes associated with altered chromatin accessibility. Detailed morphological analyses further suggest that irregularities in cell shape perpendicular to flow may alter flow sensing. Furthermore, we demonstrate the potential to ameliorate these aberrant responses using Sp1 inhibition and base editing approaches. Together, these findings identify defective endothelial adaptation to flow as a key mechanism contributing to vascular pathology in SWS and other flow-associated vascular disorders.

## Results

### Establishment of a SWS-specific pluripotent stem cell model via genome editing

The somatic nature of the causative mutation in Sturge–Weber syndrome (SWS) limits access to affected tissues, such as brain and facial vasculature, for deriving patient-specific pluripotent stem cells. To generate an isogenic disease model, we introduced the *GNAQ* c.548G>A (p.R183Q) mutation into wild-type (WT) iPSCs (409B2) using CRISPR–Cas9-mediated homology-directed repair (HDR) (Extended Data Fig. 1a–c). To enable scarless genome editing, the donor construct incorporated microhomology sequences, allowing subsequent removal of the selection cassette via microhomology-mediated end joining (MMEJ)^17^. The mutation was introduced in a heterozygous manner, and an additional synonymous substitution was included to prevent re-cleavage and enable allele-specific analysis (Extended Data Fig. 1d). Using this two-step strategy, we established SWS-specific isogenic iPSC lines, providing a robust platform for comparative phenotypic analyses.

### Development of a perfusable vessel-on-a-chip system

While GNAQ signaling has been implicated in endothelial responses to flow^15^, the behavior of mutant GNAQ under perfusion remains unclear. To address this, we developed a perfusable vessel-on-a-chip system to model flow-dependent vascular responses. The microfluidic device, composed of polydimethylsiloxane (PDMS) with parallel channels separated by micropillars, supported the formation of three-dimensional vascular networks within an extracellular matrix (Fig. 1a–d)^18^. Multiple EC types, including iPSC-derived ECs, self-organized into lumenized and interconnected vascular structures within approximately seven days, as confirmed by confocal imaging (Fig. 1a–d). Perfusion was achieved by introducing medium flow through adjacent channels, generating stable and periodic flow conditions within the vascular network (Fig. 1e–i). This platform enabled reproducible perfusion culture in a standard incubator. Using this system, we established an experimental pipeline in which WT and SWS-specific iPSC-derived ECs were differentiated and subjected to perfusion-based vascular assays (Fig. 2a).

**Fig. 1:**
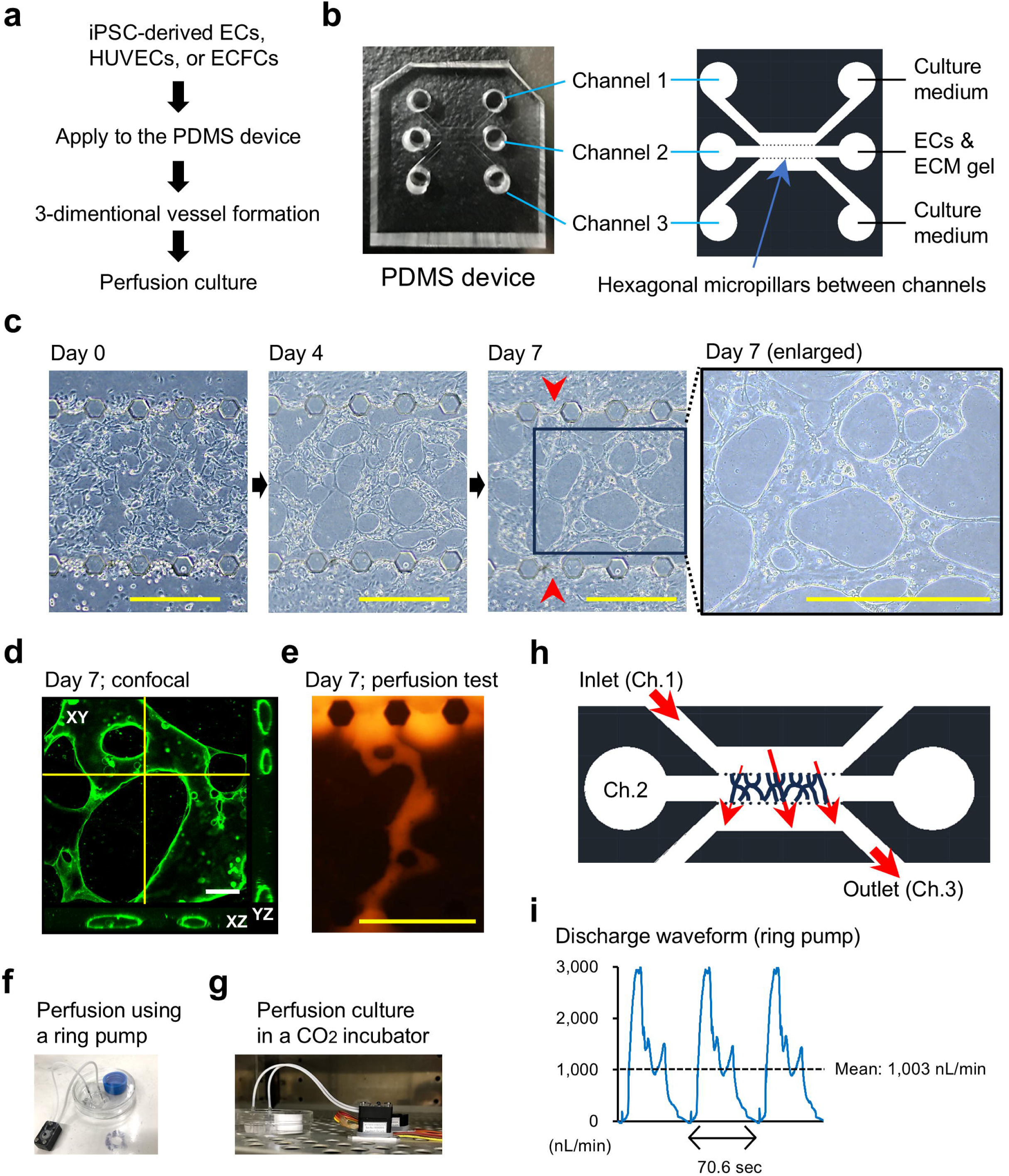
Establishment of perfusable vessel-on-a-chip. **a**, Flow of vessel-on-a-chip experiment using iPSC-derived ECs, HUVECs, or ECFCs. **b**, The PDMS device and schematic diagram. A mixture of ECs and ECM was applied to channel (Ch.) 2, and culture medium was added to Ch.1 and Ch.3. Each channel was separated by hexagonal slits. **c**, Representative phase-contrast images of ECFC-derived vessel formation in Ch.2 on days 0, 4 and 7. The formed vessels opened into the upper and lower lateral channels (red arrowheads), enabling luminal perfusion. **d**, Confocal image of 3-dimentional vessel on day 7. Green; FITC-lectin. Orthogonal cross-sectional views along the indicated yellow lines are shown. **e**, Perfusion test of vessel on day 7. Red; Rhodamine dextran. **f**, Vessel perfusion using a ring pump. Flow was generated by driving medium from the inlet (Ch.1) and withdrawing it at the outlet (Ch.3), enabling diagonal perfusion through the vessel network. **g**, Vessel perfusion culture in a CO_2_ incubator. **h**, Schematic diagram of vessel perfusion culture. Red arrows indicate the direction of medium flow, driven from Ch.1 and withdrawn at Ch.3. This configuration generated heterogeneous shear stress across the vessel network. **i**, Discharge waveform of the ring pump measured under the experimental settings in a pump-only setup (without the vessel-on-a-chip), representing the input flow profile. Scale bars represent 500 mm (yellow) and 50 mm (white). iPSC, induced pluripotent stem cell. EC, endothelial cell. HUVEC, human umbilical vein endothelial cell. ECFC, endothelial colony-forming cell. PDMS, polydimethylsiloxane. ECM, extracellular matrix.

**Fig. 2:**
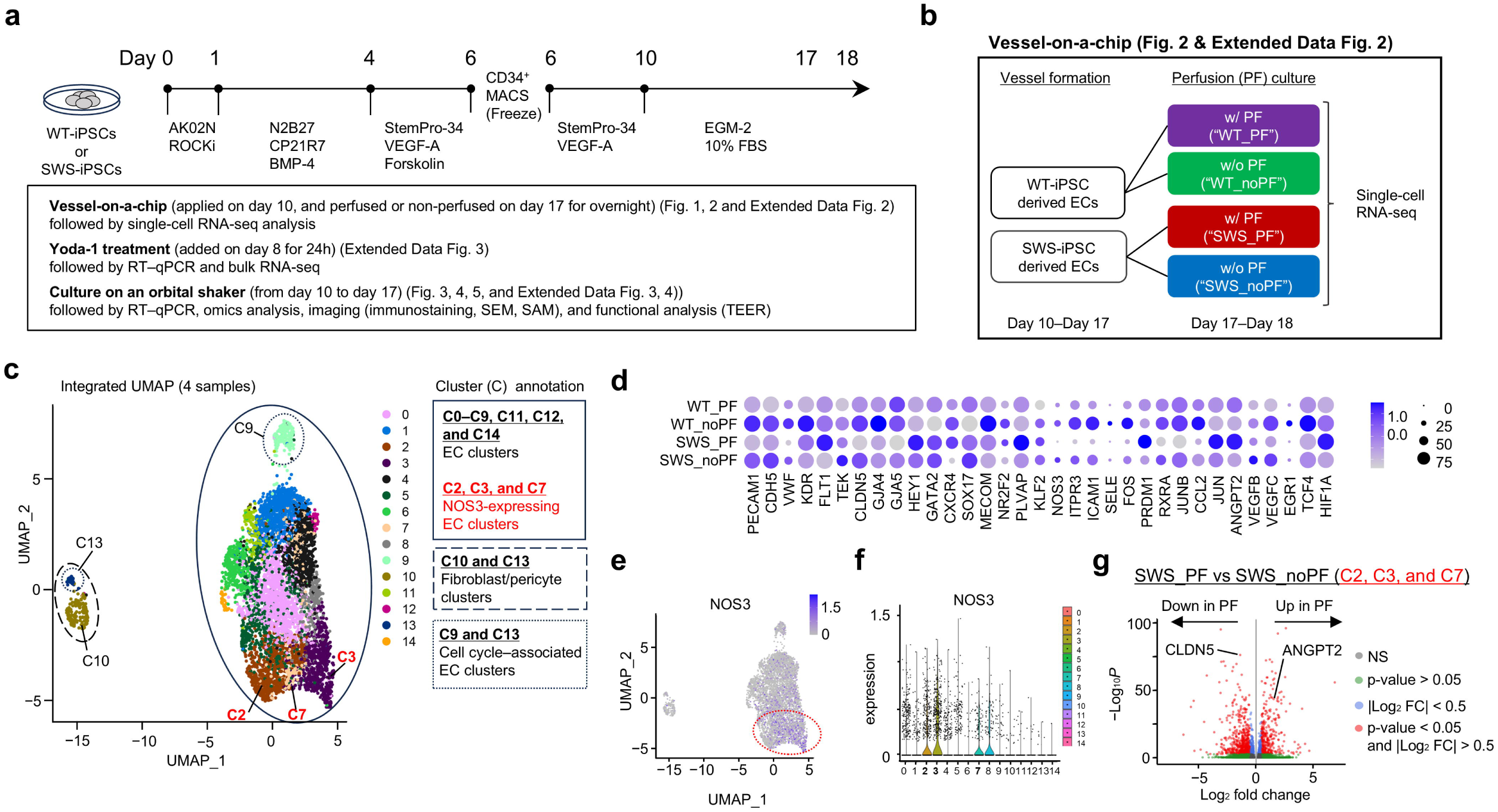
Single-cell transcriptomic analysis of iPSC-derived endothelial cells under perfusion conditions. **a**, Schematic overview of the experimental workflow and study design for functional assays. Detailed procedures are described in the Methods section. **b**, Schematic of the vessel-on-a-chip experiment. Human iPSC-derived ECs from WT or SWS were subjected to vessel-on-a-chip culture for 7 days. Perfusion (PF) was applied from day 17 for overnight culture, generating four conditions: WT with PF (WT_PF), WT without PF (WT_noPF), SWS with PF (SWS_PF), and SWS without PF (SWS_noPF). **c**, Integrated UMAP visualization of single-cell RNA-seq data from all four conditions, showing clustering of 15 populations. Clusters (C) were annotated based on gene expression profiles. **d**, Dot plots showing expression patterns of representative endothelial, shear stress–responsive, and inflammatory genes across clusters. **e**, UMAP feature plots showing EC subclusters expressing NOS3 (C2, C3, and C7). **f**, Violin plots showing NOS3 expression distribution across clusters. **g**, Differential gene expression analysis between SWS_PF and SWS_noPF conditions in NOS3-positive EC clusters (C2, C3, and C7). Volcano plot highlights significantly upregulated and downregulated genes under perfusion, including ANGPT2 and CLDN5. WT, wild-type. SWS, Sturge–Weber syndrome. PF, perfusion. UMAP, uniform manifold approximation and projection. NS, non-significant. FC, fold change.

### Single-cell transcriptomic profiling reveals altered flow responsiveness in SWS ECs

Due to substantial heterogeneity in vessel diameter within the on-chip vascular network, the wall shear stress generated by perfusion is inherently variable. To resolve perfusion-dependent effects at single-cell resolution, we performed single-cell RNA-sequencing (scRNA-seq) on vascular cells derived from WT and SWS conditions, with or without perfusion (PF), yielding four experimental groups (Fig. 2a,b). Integrated UMAP analysis identified 15 distinct clusters (Fig. 2c). Based on canonical marker expression, these clusters were annotated as (i) vessel-forming ECs (C0–C9, C11, C12, and C14), (ii) cell cycle–associated ECs (C9 and C13), and (iii) pericytes or fibroblasts (C10 and C13) (Fig. 2c and Extended Data Fig. 2a,b). We next examined transcriptional responses to perfusion. While endothelial identity was preserved across all conditions, perfusion induced broad transcriptional remodeling, with distinct response patterns observed between WT and SWS cells (Fig. 2d).

Focusing on flow-responsive endothelial states, we identified subclusters (C2, C3, and C7) showing modest but relatively higher expression of NOS3 (endothelial nitric oxide synthase, eNOS) (Fig. 2e,f), a feature characteristic of quiescent endothelial states under laminar flow exposure^19–21^. These clusters were positioned opposite the proliferative EC clusters along the UMAP_2 axis, suggesting that this transcriptional axis reflects the balance between proliferative and flow-adapted quiescent endothelial states under laminar-like perfusion conditions. To define SWS-specific responses to perfusion, we performed differential expression analysis within these NOS3-enriched clusters. Notably, under perfused conditions, SWS ECs exhibited upregulation of ANGPT2 (angiopoietin-2) and downregulation of CLDN5 (claudin-5) (Fig. 2g and Extended Data Fig. 2c,d). ANGPT2 is a key regulator of endothelial activation and vascular remodeling^22^ and has been reported to be elevated in SWS-associated vasculature^23^. In contrast, CLDN5 encodes a tight junction protein essential for endothelial barrier integrity, particularly in the blood–brain barrier^24–26^. Together, these alterations suggest a shift toward a destabilized endothelial state with impaired barrier function in SWS under perfusion. In line with these changes, over-representation analysis^27^ revealed significant downregulation of the Notch signaling pathway in perfused SWS ECs (Extended Data Fig. 2e). Given the established role of Notch signaling in arterial specification and endothelial stability^28,29^, this suppression further supports a shift toward a more activated endothelial state accompanied by reduced stability and impaired maturation in SWS ECs under perfused conditions. Collectively, these findings indicate that SWS ECs exhibit aberrant transcriptional responses to perfusion, characterized by endothelial activation, impaired barrier integrity, and disrupted maturation programs, suggesting defective adaptation to flow-responsive endothelial states.

We next assessed whether intrinsic cellular properties differed between WT and SWS conditions. Independent of perfusion status, SWS samples showed a relative reduction in flow-responsive EC clusters (C2, C3, and C7) and a concomitant increase in proliferative populations, including cell cycle–associated ECs (C9) and cycling perivascular cells (C13) (Extended Data Fig. 2f). Consistent with this observation, SWS-derived iPSCs exhibited significantly increased proliferative capacity compared with WT controls, as demonstrated by CFSE analysis (Extended Data Fig. 2g). These results indicate that the SWS-associated mutation confers a baseline proliferative bias.

Importantly, this intrinsic proliferative tendency does not account for the perfusion-dependent transcriptional alterations described above, indicating that altered responsiveness to perfusion represents a distinct and defining feature of SWS ECs.

### High-dose Yoda-1 partially recapitulates flow-induced changes

To assess whether chemical activation of mechanotransduction could mimic flow responses, we treated ECs with the Piezo agonist Yoda-1 (Extended Data Fig. 3a)^30,31^. Yoda-1 induced dose-dependent upregulation of ANGPT2 and downregulation of CLDN5, partially recapitulating the scRNA-seq findings (Extended Data Fig. 3b,c). Transcriptomic analysis revealed distinct transcriptional responses to Yoda-1 between WT and SWS ECs, with greater divergence observed at higher Yoda-1 concentrations (32 μM; hereafter referred to as the treatment (Tx) condition) (Extended Data Fig. 3d). In SWS ECs, the Tx condition was associated with activation of TNFα and mTORC1 signaling pathways (Extended Data Fig. 3e). However, CLDN5 downregulation was observed only under the Tx condition (Extended Data Fig. 3c), raising the possibility that at least part of these changes reflected cytotoxicity associated with high-dose Yoda-1 (32 μM).

### Simple laminar pulsatile flow reveals morphological abnormalities in SWS ECs

We next applied a simplified laminar flow model using an orbital shaker system, in which cells at the periphery are exposed to pulsatile shear stress (Fig. 3a)^32,33^. Under these conditions, WT ECs exhibited elongation and alignment along the flow direction, whereas SWS ECs displayed irregular cell borders and prominent spiky surface structures (Fig. 3b). Consistent with the on-chip scRNA-seq results, CLDN5 expression was reduced in SWS ECs (Fig. 3c,d). Ultrastructural analysis by SEM revealed minimal intercellular gaps in WT cells, while SWS ECs exhibited widened intercellular spaces and protrusive structures reminiscent of endothelial filopodia^34^ (Fig. 3e). To further characterize these morphological differences in three dimensions, we performed scanning acoustic microscopy (SAM)-based topographic analysis^35,36^. In the peripheral regions exposed to shear stress, both groups showed reduced surface roughness compared to the central regions; however, SWS ECs remained significantly rougher than WT ECs (Fig. 3f–j and Extended Data Fig. 3f,g). Cross-sectional analysis along the flow direction revealed distinct differences in height profiles between WT and SWS ECs (Fig. 3g). To dissect the origin of these differences, we applied Gaussian mixture model (GMM) fitting^37^ to estimate nuclear and cytoplasmic height distributions (Fig. 3k–m and Extended Data Fig. 3h–j). While shear stress reduced the nucleus–cytoplasm height differences in both groups, this reduction was significantly attenuated in SWS ECs compared with WT ECs specifically in the peripheral regions. Importantly, these differences were independent of cellular density (Extended Data Fig. 3k). Together, these findings indicate that SWS ECs exhibit altered morphological adaptation to shear stress, particularly in vertical cellular architecture along the flow direction.

**Fig. 3:**
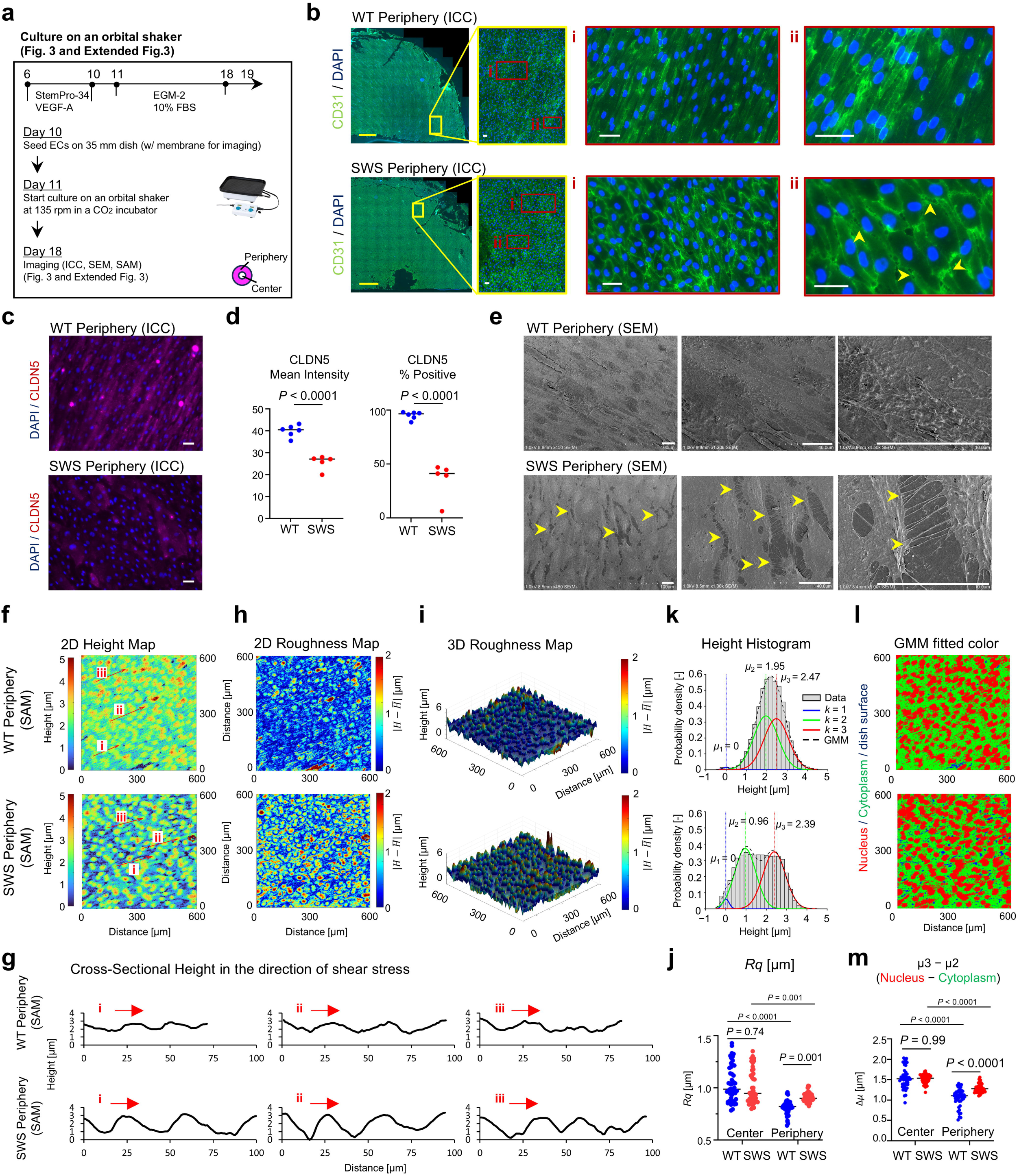
Shear stress induces morphological and height alterations in SWS endothelial cells. **a**, Experimental workflow for orbital shear stress culture. iPSC-derived ECs were seeded at day 10 and cultured under orbital shaking (135 rpm) from day 11 to day 18, followed by imaging. **b**–**d**, ICC of WT and SWS ECs at the dish periphery. CD31 (**b**; green) and CLDN5 (**c**; red) with DAPI staining were shown. SWS cells exhibited irregular, spiky membrane structures (arrowheads). CLDN5 intensity and CLDN5-positive cells were quantified (**d**) and statistical significance was determined by an unpaired two-tailed Student’s t-test (WT: n = 6, SWS: n = 5). Lines indicate the median. Yellow scale bars, 2 mm; white scale bars, 50 μm. **e**, SEM images of ECs at the dish periphery. SWS cells displayed intercellular gaps and filopodia-like protrusions (arrowheads). Scale bars, 20 μm. **f**, **g**, SAM-based height maps (**f**) and corresponding cross-sectional profiles (**g**). Cross-sectional analysis along the flow direction revealed distinct height profiles between WT and SWS ECs. **h**, **i**, 2D (**h**) and 3D (**i**) roughness maps. **j**, Quantification of surface roughness (Rq) at the center and periphery of the culture dish. Lines indicate the median. **k**–**m**, Height distributions analyzed by GMM fitting (**k**), identifying three components corresponding to dish (*k* = 1; μ1), cytoplasm (*k* = 2; μ2), and nucleus (*k* = 3; μ3) (**l**). The nuclear–cytoplasmic height difference (μ3 − μ2) is quantified (**m**). In panels **f**, **h**, **i**, **k**, and **l**, representative images from the peripheral region of WT and SWS cultures are shown in the upper and lower rows, respectively. Statistical significance (**j**, **m**) was evaluated by two-way ANOVA followed by Šídák’s multiple comparisons test. Lines indicate the median. ICC, immunocytochemistry. SEM, scanning electron microscopy. SAM, scanning acoustic microscopy. GMM, gaussian mixture model.

### Flow significantly modulates endothelial barrier function in SWS ECs

Morphological adaptation to shear stress is thought to influence endothelial integrity. To enrich for cells responsive to laminar flow, centrally located cells were selectively removed prior to perfusion, and transendothelial electrical resistance (TEER) was measured immediately after passaging (Fig. 4a). Physiologically relevant laminar shear stress is known to enhance endothelial barrier function in both monolayer and engineered vessel systems^3,4,38^. Consistently, using cellZScope measurements^39^, WT cells pre-exposed to perfusion exhibited increased TEER after passaging, whereas SWS cells failed to show such enhancement (Extended Data Fig. 4a). This result was further validated using the Maestro Z platform, which enables multi-frequency impedance measurements^40^, where perfusion increased TEER (1 kHz resistance) in WT cells but not in SWS cells; instead, TEER was reduced in SWS cells following perfusion (Fig. 4b), indicating impaired barrier reinforcement. In contrast, high-frequency resistance (41.5 kHz), a surrogate for cell coverage of the well, remained within a similar range across WT and SWS conditions (Fig. 4c), suggesting that the observed differences were not attributable to changes in cellular density. Consistent with these findings, the Barrier Index, calculated as a ratio of low- to high-frequency resistance measurements, was significantly decreased in SWS under perfusion (Fig. 4d). This phenotype was robust across time points and seeding densities (Extended Data Fig. 4b). Together, these results demonstrate that SWS ECs fail to reinforce barrier function in response to shear stress and allow barrier-deficient ECs to persist under flow, leading to their progressive accumulation under perfusion.

**Fig. 4:**
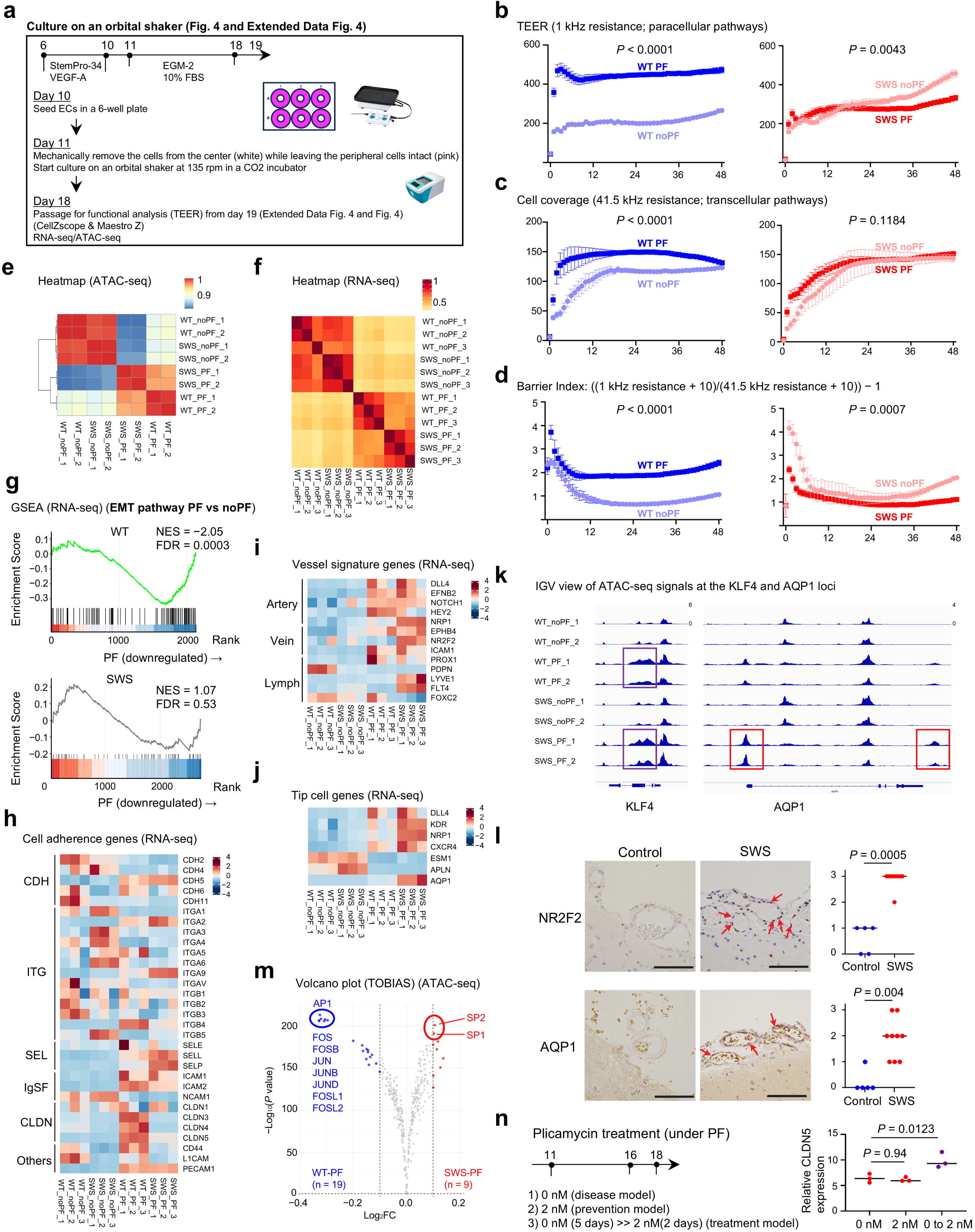
Laminar pulsatile flow induces aberrant barrier function and transcriptional reprogramming in SWS endothelial cells. **a**, Experimental design for orbital shear stress culture and functional assays. ECs at the peripheral region exposed to flow were enriched by mechanical removal of central cells prior to PF, followed by passaging and TEER measurement. **b**, TEER measurements (1 kHz resistance; paracellular pathway) using Maestro Z. PF enhanced barrier function in WT ECs, whereas SWS ECs failed to exhibit this response and instead showed decreased TEER following PF. **c**, Cell coverage assessed by high-frequency resistance (41.5 kHz; transcellular pathway), showing comparable viability between WT and SWS ECs. **d**, Barrier index calculated from 1 kHz and 41.5 kHz resistance measurements, demonstrating impaired barrier integrity in SWS ECs under PF. Data **b**–**d** were presented as mean ± SD and were analyzed by two-way repeated measures ANOVA to assess the effects of genotype and time, as well as their interaction. **e**, **f**, Heatmaps of ATAC-seq (e) and RNA-seq (f) data showing global transcriptional and chromatin accessibility changes. Samples were primarily segregated by PF exposure, with increased divergence between WT and SWS under PF conditions. **g**, GSEA highlighting pathway specifically altered in SWS ECs under PF condition. **h**, Expression of cell adhesion–related genes from RNA-seq data. SWS ECs under PF exhibited reduced expression of tight junction components (e.g., claudins) and increased expression of adhesion molecules such as selectins and integrins. **i**, Expression of vessel signature genes, showing some venous and lymphatic genes upregulated in SWS ECs under PF. **j**, Expression of tip cell–associated genes, showing selective upregulation in SWS ECs under PF. **k**, Representative IGV tracks of chromatin accessibility showing consistently open chromatin at the KLF4 locus in PF conditions in both WT and SWS ECs, whereas increased accessibility at the AQP1 locus is observed selectively in SWS-PF ECs. **l**, Validation in patient tissue samples. Expression of venous- and tip cell-associated genes was increased in SWS lesions compared to controls. Statistical significance was determined by the Mann-Whitney U test for non-parametric analysis of scoring data. Lines indicate the median. **m**, Transcription factor motif accessibility analyzed by TOBIAS on ATAC-seq data. SP-family motifs (SP1/2) showed increased accessibility in SWS ECs under PF, whereas accessibility of AP1 motifs was globally reduced in SWS under the same condition. **n**, Effects of SP1 inhibition (Plicamycin) under PF conditions. Treatment initiated at day 5 (treatment model) resulted in greater improvement in barrier function compared to treatment from day 1 (prevention model). Statistical significance was determined by one-way ANOVA followed by Dunnett’s multiple comparisons test (n = 3 per group). Lines indicate the median. TEER, transendothelial electrical resistance. IGV, Integrative Genomics Viewer.

### Flow induced global transcriptomic changes and chromatin status alterations

To investigate the causes of these morphological and functional changes, bulk RNA-seq (n = 3 in each group) and ATAC-seq (n = 2 in each group) analyses were performed in parallel. The four sample types were broadly classified based on the presence or absence of PF in the heatmap. However, a comparison between WT and SWS revealed that the differences between the PF groups were more pronounced than those between the non-PF groups (Fig. 4e,f). PCA also indicated a greater separation between the PF groups (Extended Data Fig. 4c,d). Chromatin accessibility at the promoter and transcription start site (TSS) regions, a feature generally associated with transcriptional activity^41^, showed an increasing trend in the SWS-PF group (Extended Data Fig. 4e). To further assess the transcriptional programs associated with PF exposure, we performed a gene set enrichment analysis (GSEA). PF induced a consistent downregulation of core cell cycle–related pathways in both WT and SWS, including E2F targets, G2M checkpoint, and MYC targets (Extended Data Fig. 4f). In contrast, the epithelial–mesenchymal transition (EMT) pathway, which has been linked to endothelial plasticity and EndoMT-related programs and is typically suppressed under physiological flow conditions^42^, exhibited divergent regulation, being suppressed in WT but not in SWS (Fig. 4g), suggesting impaired maintenance of endothelial integrity in SWS. Reflecting these alterations in the EMT pathway, we next examined the expression of cell adhesion–related genes. The SWS-PF group showed decreased expression of claudins, including claudin 5, alongside increased expression of selectins involved in leukocyte–endothelial adhesion and elevated integrin α9 expression, which is associated with lymphangiogenesis (Fig. 4h). Examination of vascular-related genes revealed that, while arterial markers remained prominent in the PF groups overall, the SWS-PF group showed elevated expression of venous (NR2F2) and lymphatic-related genes (LYVE1 and FLT4) (Fig. 4i). Furthermore, while stalk cell–associated genes were commonly upregulated in the PF groups (Extended Data Fig. 4g), certain tip cell–associated genes^43,44^, including AQP1, a recently described marker associated with endothelial angiogenic sprouting^45^, were specifically upregulated in the SWS-PF group (Fig. 4j). In the chromatin accessibility data derived from ATAC-seq, we observed that the stalk cell–associated gene KLF4 showed consistently open chromatin in the PF groups, whereas the tip cell–associated gene AQP1 and the lymphatic-associated gene FLT4 exhibited increased accessibility specifically in the SWS-PF group (Fig. 4k and Extended Data Fig. 4h). To determine whether this phenomenon could also be observed in patient tissue samples, we examined the expression of NR2F2, AQP1, and SOX18 in the abnormally dilated capillary-like blood vessels of the cerebral leptomeninges. NR2F2 and AQP1 expression was significantly increased, with a non-significant increase in SOX18, a transcription factor implicated in lymphatic development^46^ (control, n = 5; SWS, n = 9) (Fig. 4l Extended Data Fig. 4j,k). As a potential underlying mechanism, comparison of chromatin accessibility between SWS-PF and WT-PF revealed increased accessibility of SP1, SP2, and SP3 binding motifs in SWS-PF (Fig. 4m and Extended Data Fig. 4i). Conversely, AP-1 transcription-factor-binding motifs exhibited reduced accessibility in the SWS group, irrespective of PF (Fig. 4m and Extended Data Fig. 4i). Consistent with this, treatment with an SP1 inhibitor (Plicamycin)^47^ during perfusion culture resulted in an increase in CLDN5 expression, indicating partial phenotypic rescue (Fig. 4n). Collectively, these findings indicate that SWS-PF ECs partially recapitulate flow-induced transcriptional programs observed in WT-PF cells, while concurrently acquiring aberrant features, including tip-like endothelial traits, upregulation of venous and lymphatic markers, and altered cell adhesion profiles. Chromatin accessibility analysis and pharmacological inhibition further implicate SP family transcription factors in driving this phenotype.

### Adenine base editing (ABE) genome editing specifically repairs disease-causing mutations and reverses the phenotype of disease cell models

Finally, given the challenge of targeting the SP1 transcription factor as a disease-specific therapeutic strategy due to its broad involvement in fundamental cellular processes and the associated risk of systemic toxicity upon its inhibition^48^, we sought to directly correct the causal mutation at the DNA level. We first evaluated conventional CRISPR–Cas9 approaches; however, no suitable PAM site compatible with SpCas9 (NGG PAM requirement) was available in proximity to the mutation, precluding precise targeting (Fig. 5a). When using guide RNAs (gRNAs) designed to cleave at the closest possible sites, a distance of 10–12 nt between the variant and the predicted Cas9 cleavage site presumably prevents allele-specific discrimination, thereby increasing the likelihood of unintended cleavage of the wild-type allele, consistent with the well-established tolerance of Cas9 to mismatches at PAM-distal regions^49,50^. To overcome this limitation, we employed adenine base editing (ABE), which enables A-to-G substitutions within a defined PAM-distal editing window^51^ (Fig. 5b) and has been successfully applied to the therapeutic correction of pathogenic variants^52^, allowing precise correction of the disease-causing mutation (c.548G>A) positioned within this window (Fig. 5c). Importantly, potential bystander edits were predicted to result only in synonymous substitutions, due to the local codon context (Fig. 5c). We systematically evaluated three gRNAs of varying lengths (truncated and extended designs) in combination with two ABE variants, ABE8.8-m and ABE7.10 (Extended Data Fig. 5a)^51,52^. Across all conditions, editing was largely restricted to the intended bases, with correction of the mutant allele and only synonymous bystander substitutions detected (Extended Data Fig. 5b). Among the gRNAs tested, the G + 19 nt gRNA exhibited the highest editing efficiency and precision across both ABE variants. Notably, under this condition, ABE8.8-m achieved a higher editing efficiency than ABE7.10 while maintaining comparable precision. Using this optimized condition (G + 19 nt gRNA with ABE8.8-m), we successfully established SWS-corrected-iPSC clones harboring precise A-to-G correction of the mutant allele, without detectable modification of the wild-type allele (Extended Data Fig. 5c). Moreover, ECs differentiated from the ABE-corrected SWS-iPSC clone exhibited restored CLDN5 expression under perfusion conditions, compared to their isogenic mutant counterparts, indicating functional rescue of endothelial barrier properties (Fig. 5d and Extended Data Fig. 5d). Collectively, these findings demonstrate that ABE-mediated correction enables precise and functional repair of the pathogenic mutation in this SWS model.

**Fig. 5:**
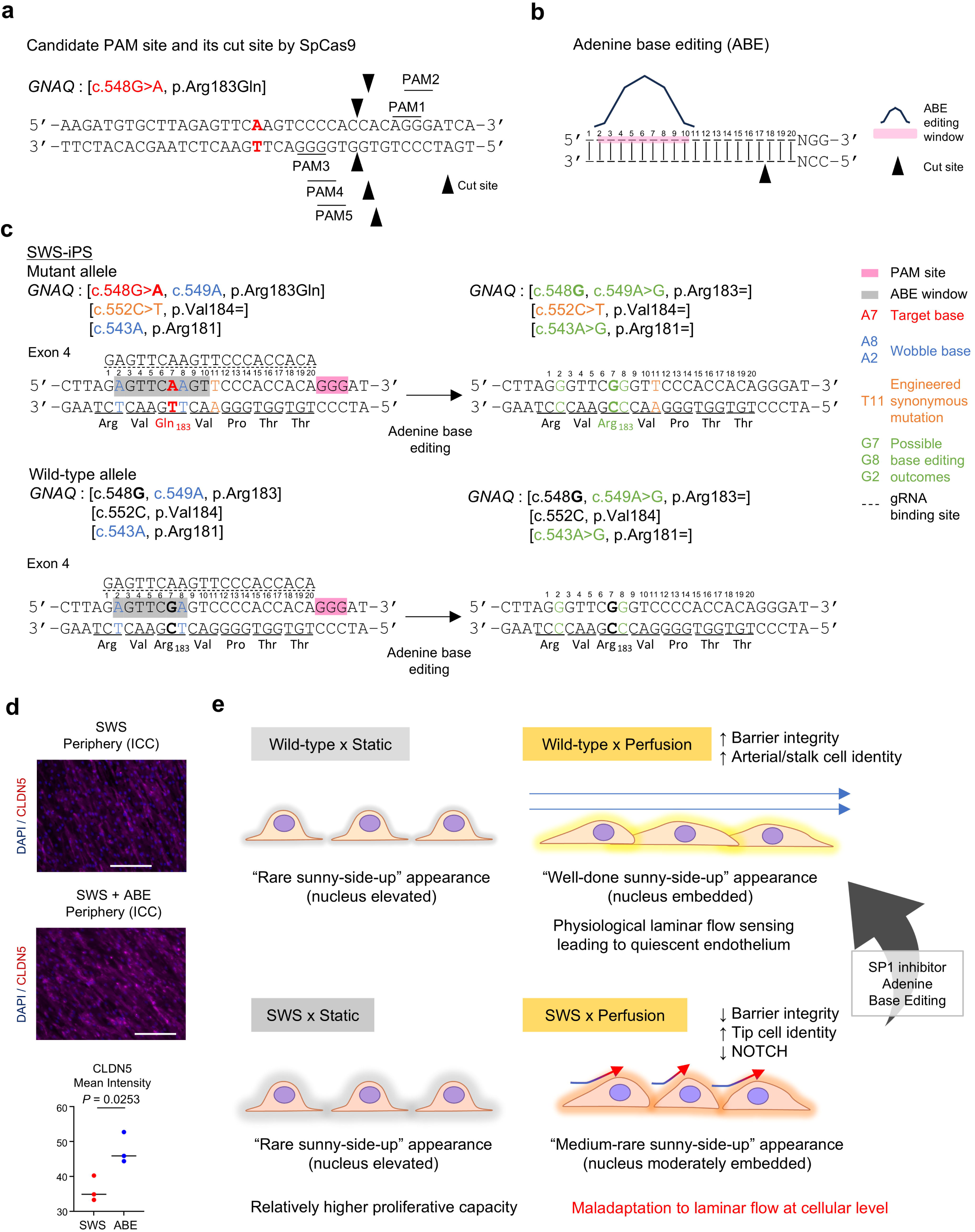
Adenine base editing enables precise correction of the disease-causing mutation and rescues endothelial phenotypes. **a**, Schematic illustrating the limitation of conventional CRISPR–Cas9-mediated genome editing for the GNAQ c.548G>A mutation. No suitable PAM-proximal gRNA could be designed to enable allele-specific targeting. **b**, Overview of adenine base editing (ABE), which induces A-to-G conversion within a defined editing window distal to the PAM site. **c**, Design of ABE targeting strategy for the GNAQ mutant allele. The pathogenic c.548G>A mutation is positioned within the editable window, enabling precise reversion to the wild-type sequence. Predicted bystander edits in both mutant and wild-type alleles result exclusively in synonymous substitutions. **d**, Immunofluorescence analysis of CLDN5 expression in SWS ECs with or without ABE correction under flow conditions. ABE-corrected ECs exhibited restoration of CLDN5 expression compared to uncorrected SWS ECs. Quantification of CLDN5-positive cells was shown. Statistical significance was determined by an unpaired two-tailed Student’s t-test (n = 3 per group). Lines indicate the median. **e**, Summary of flow-mediated SWS vascular cellular model pathogenesis and possible treatment approach including SP1 inhibitor or ABE. gRNA, guide RNA. ABE, adenine base editing. EC, endothelial cell. ICC, immunocytochemistry.

## Discussion

Our study demonstrates that perfusion elicits aberrant responses in vascular ECs derived from iPS cells harboring the SWS mutation, implicating flow maladaptation as a key contributor to disease pathology. In this context, our findings further suggest that structural abnormalities at the cell surface may underlie this altered mechanosensory response (Fig. 5e).

A key strength of this study lies in the use of disease-model iPS cells in which a single mutant allele was heterozygously introduced, thereby avoiding the confounding effects associated with conventional overexpression systems. To enable precise phenotypic comparisons, we employed isogenic iPS cells generated by introducing the mutation into wild-type cells, thereby minimizing variability arising from differences in the genetic background, a known limitation in patient-derived iPS cell models^53–55^. In addition, our seamless genome editing strategy based on MMEJ, which avoids the introduction of exogenous sequences, enhances the fidelity of genotype–phenotype relationships.

In vivo, blood vessels are three-dimensional and highly heterogeneous tissues that are continuously exposed to mechanical forces, including cyclic stretch from blood pressure and shear stress from blood flow^4,20^. In this study, we reconstructed perfusable capillary-like vascular networks in three dimensions using an on-chip system (Fig. 1). Establishing conditions under which iPSC-derived ECs could reproducibly form stable and perfusable three-dimensional vascular networks, rather than simple two-dimensional monolayers, was a major technical prerequisite for this study. In our previous study, only a limited number of ECs could form perfusable vascular networks^18^. Under optimized conditions for matrix composition, culture medium, cell-seeding density, and culture timing, both WT and SWS iPSC-derived ECs self-organized into interconnected and lumenized vascular networks that supported continuous perfusion. Unlike large vessels such as arteries, arterioles, and veins, which possess well-defined three-layered structures, SWS is characterized by malformed vessels resembling capillary-to-venous structures^11^, rendering our experimental system particularly relevant to the disease context. To capture the complexity of endothelial responses to shear stress, we applied scRNA-seq to the on-chip model. At the same time, to dissect causal relationships between perfusion and disease phenotypes, reductionist experimental systems are also required. We therefore employed an orbital shaker to generate laminar pulsatile flow in a simplified setting, enabling the analysis of morphological, transcriptional, and chromatin changes. Together, these approaches demonstrate that key phenotypes observed in the complex on-chip system are reproducible under simplified flow conditions, allowing for a more detailed interrogation of the underlying morphological and molecular mechanisms. Therapeutic strategies for SWS have increasingly focused on targeting enhanced Gq signaling, its downstream effector PLCβ, and related pathways such as ANGPT2, with emerging evidence of efficacy^23^. These studies have provided important insights into the signaling mechanisms underlying disease pathology. Building on this framework, our findings further point to dysregulation at the transcriptional network level, characterized by increased SP1 activity and reduced AP-1 activity. These alterations appear to arise from the altered ability of SWS ECs to appropriately sense and respond to laminar flow stimuli, which, in normal ECs, would induce the acquisition of arterial or stalk-cell phenotypes. Targeting such transcriptional programs may therefore represent an additional avenue for correcting disease-associated endothelial dysfunction. Recent studies by Bischoff et al. have highlighted the importance of perfusion in regulating leukocyte migration^56^ and EC alignment^57^ in SWS. Extending these observations, our study provides a mechanistic link between aberrant flow sensing and downstream transcriptional rewiring, thereby refining the current understanding of how perfusion contributes to disease pathology.

Previous studies have shown that chromatin accessibility at AP-1 binding motifs increases under disturbed flow conditions^58^; similarly, AP-1 expression is enhanced under physiological laminar pulsatile flow^59^. Together, these findings highlight the dynamic responsiveness of this transcription factor to hemodynamic cues. In the present study, we observed reduced chromatin accessibility at AP-1 binding motifs in SWS ECs under perfusion. The flow conditions generated by the orbital shaker differ from the laminar or disturbed flow patterns produced by well-controlled microfluidic platforms (e.g., ibidi), which may partly account for this discrepancy. In this context, the altered behavior of AP-1 observed here may reflect an abnormal response to flow stimuli in SWS ECs under the experimental conditions used. Given the central role of flow-responsive transcriptional networks in vivo, these findings suggest that this regulation may be particularly sensitive to both mechanical and disease-associated contexts.

Although the applicability of adenine base editing (ABE) depends on the mutation context, this approach offers the potential for precise genetic correction in diseases such as SWS. In the present study, we identified ABE configurations that minimize bystander editing, thereby supporting the feasibility of targeted therapeutic interventions. It should be noted that the guide RNA used here was designed to recognize the mutant allele together with a linked synonymous variant, and further optimization of mutation-selective guide RNAs will be important for therapeutic applications.

Previous studies have demonstrated that nanoscale surface structures, such as the endothelial glycocalyx, play critical roles in mechanosensing and the transduction of shear stress^38,60^. At the opposite end of the spatial spectrum, macroscale hemodynamic disturbances, including flow separation and turbulence at the vessel level, have been implicated in vascular pathologies such as atherosclerosis^3,4^. Together, these studies highlight the importance of structural features across multiple scales in shaping endothelial responses to hemodynamic cues. In contrast, the present study identifies a previously underappreciated intermediate scale, showing that abnormalities at the level of individual ECs—arising from altered three-dimensional cellular morphology, including cytoplasmic and nuclear architecture—may perturb local flow profiles and impair appropriate responses to laminar shear stress (Fig. 5e). This multiscale perspective expands the conceptual framework linking cellular morphology to endothelial mechanobiology. Importantly, this perspective extends beyond therapeutic strategies focused primarily on individual molecular pathways, suggesting that the restoration of cellular morphology and surface architecture may provide a complementary framework for understanding and potentially treating not only vascular malformations but also a broader spectrum of vascular diseases.

## Acknowledgements

We are grateful to Mervin C. Yoder for his continued mentorship throughout this work. We thank Hiromi Misawa for administrative assistance and Nara Medical University Research Core for access to core facilities and equipment. We also thank Naotaka Nakazawa (Kindai University) for insightful comments and creative suggestions that greatly improved this study. This work was supported by JST FOREST (JPMJFR235D to K.B.); the Japan Agency for Medical Research and Development (AMED; JP22ek0109605 and JP26ym0126809 to K.B.); JSPS KAKENHI (JP21K15909 and JP24K02426 to K.B.); the 2020 iPS Academia Japan Grant (to K.B.); the 2020 Medical Research Grant from the Takeda Science Foundation (to K.B.); the 2020 Ichiro Kanehara Foundation for the Promotion of Medical Sciences and Medical Care (to K.B.); the 2021 Research Grant from the Japan Epilepsy Research Foundation (to K.B.); and the 2023 Research Grant from the Mother and Child Health Foundation (to K.B.).

This work was also supported by the Advanced Research Infrastructure for Materials and Nanotechnology in Japan (ARIM), Ministry of Education, Culture, Sports, Science and Technology (MEXT; Proposal No. JPMXP1222KT1334). We acknowledge the NGS Core Facility at the Research Institute for Microbial Diseases, The University of Osaka, for sequencing and data analysis, and the CiRA Foundation (CiRA_F, Kyoto, Japan) for single-cell RNA-sequencing and technical support.

## Author Contributions

Conceptualization, K.B., R.Y., Y.U., K.H.; Methodology, K.B., R.Y., J.M., J.Y., Y.K., Y.O., K.S., K.W., T.H., Y.U., K.H.; Investigation, K.B., J.M., K.M., S.W., Y.K., A.Shirono, T.H., Y.U., K.H.; Resources, R.Y., A.K., M.T., M.I., A.Shindo; Writing, K.B., R.Y., J.Y., K.S., K.W., Y.U., K.H.; Supervision, K.B., K.H.; Funding acquisition, K.B., K.Y., and K.H.

## Declaration of generative AI

During manuscript preparation, the authors used AI-assisted tools solely for language editing and grammar improvement. All scientific content, interpretation, and conclusions were generated and verified by the authors, who take full responsibility for the manuscript.

## Data Availability

scRNA-seq data are available in the Gene Expression Omnibus (GEO) under accession number GSE330493. Bulk RNA-seq data are available under GSE330654 (Yoda-1) and GSE330603 (orbital shaker). ATAC-seq data are available under GSE330603.

Source data for all main and Extended Data figures are provided with this manuscript. Plasmids, iPSC lines, and other materials generated in this study are available from the corresponding author upon reasonable request.

Any additional information required to reanalyze the data reported in this paper is available from the corresponding author upon reasonable request.

## Extended Data Figure Legends

**Extended Data Fig. 1.**
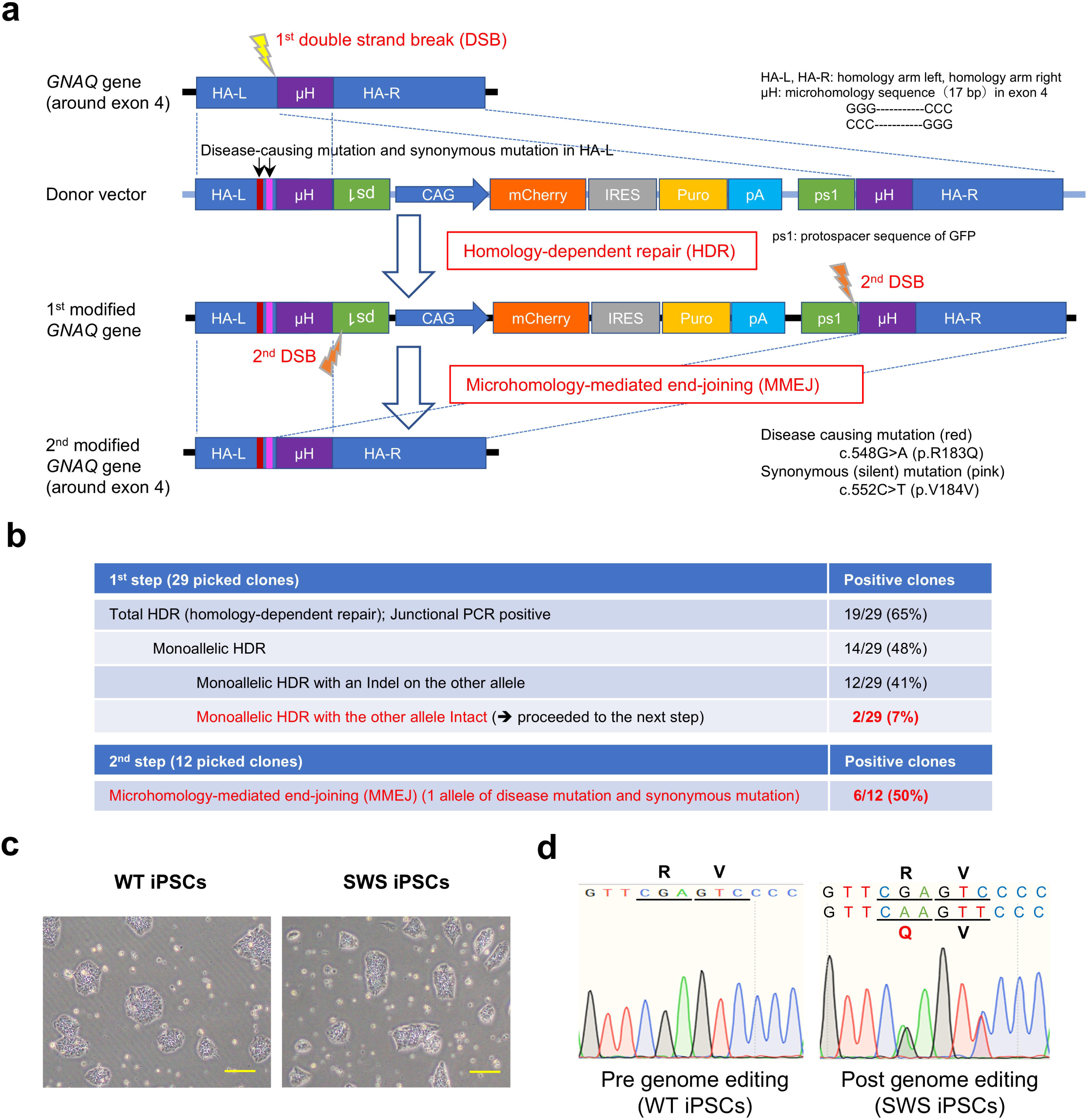
Derivation of Sturge-Weber syndrome-specific iPSCs by genome editing using CRISPR–Cas9. **a**, Schematic overview depicting the genomic *GNAQ* locus (around exon 4), donor vector, and sequentially modified *GNAQ* gene. Left and right homology arms overlap, generating a 17 bp tandem microhomology sequence (mH, purple) flanking the selection cassette. The disease-causing mutation (c.548G>A, red) and the synonymous (silent) mutation (c.552C>T, pink) were in the left homology arm (HA-L). The selection cassette was flanked by a pair of inversely oriented EGFP-derived protospacers (ps1, green) within the tandem repeat of mH. *GNAQ* gene targeting used the CRISPR–Cas9 (yellow bolt, 1^st^ double-strand break (DSB)), leading to homology-dependent repair (HDR) with the donor vector. Upon transfection of targeted clones with the second CRISPR–Cas9 (orange bolts, 2^nd^ DSB), DSBs were generated flanking the cassette, proximal to the engineered mH. Repair by microhomology-mediated end-joining (MMEJ) scarlessly excised the cassette, resulting in the desired editing outcome (2^nd^ modified *GNAQ* gene). **b**, Summary table of the genome editing experiments. In the 1^st^ step, monoallelic HDR was achieved in 14/29 clones (48%), of which 2/29 (7%) retained the other allele intact. After the 2^nd^ step, the efficiency of microhomology-mediated repair was 6/12 (50%). **c**, Representative phase-contrast images of iPSCs before and after the genome editing. Both clones maintained the typical morphology of undifferentiated iPSCs. **d**, Representative Sanger sequencing chromatograms confirming the presence of the disease mutation and synonymous mutation in the genome-edited iPSC clone (SWS-iPSCs). Scale bars represent 200 mm. DSB, double-strand break. HDR, homology-dependent repair. MMEJ, microhomology-mediated end-joining. WT, wild-type. SWS, Sturge–Weber syndrome.

**Extended Data Fig. 2.**
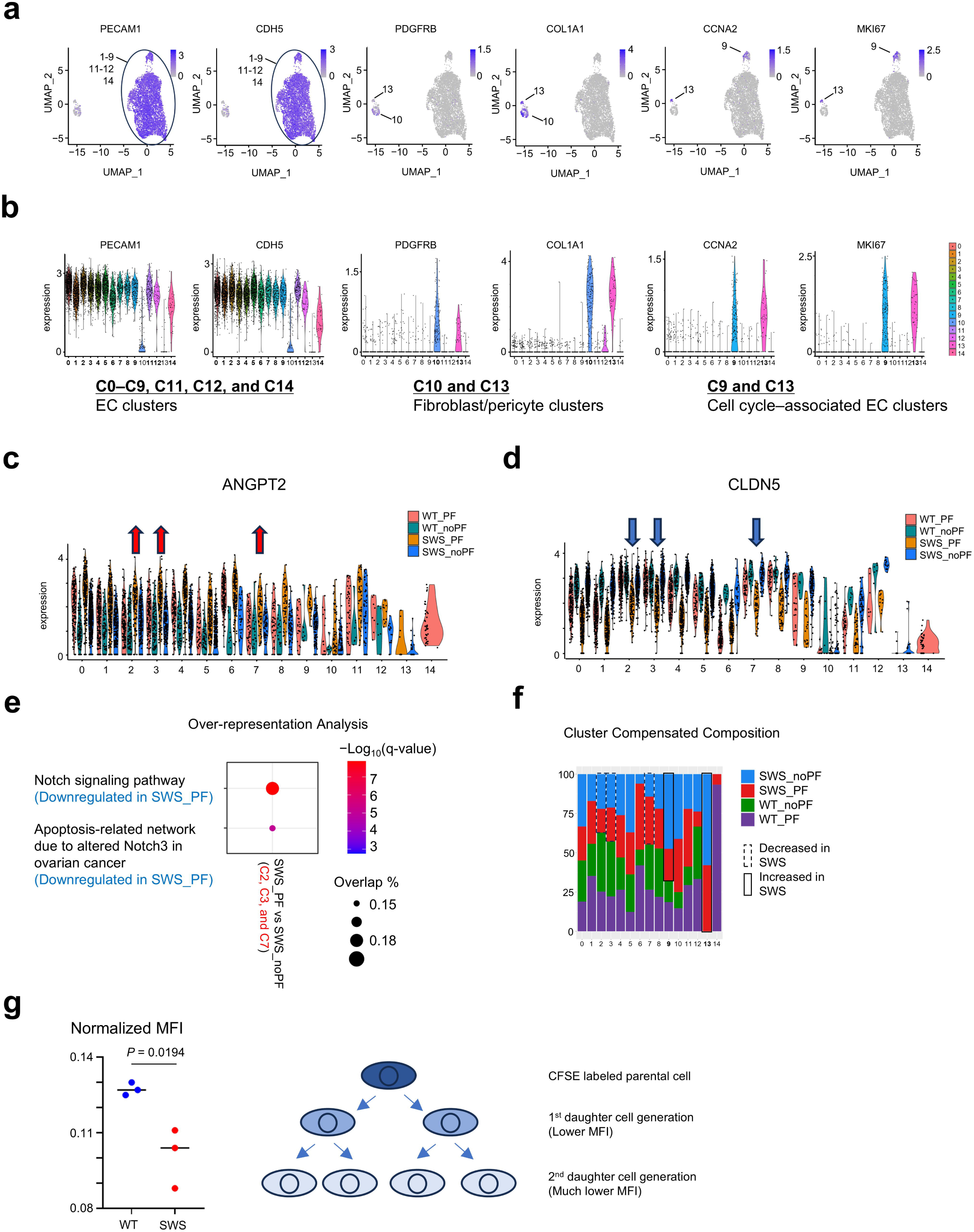
Characterization of cell clusters and pathway enrichment analysis in iPSC-derived endothelial cells. **a**, UMAP feature plots showing expression of EC genes (PECAM1, CDH5) (in C0–C9, C11, C12 and C14), mesenchymal genes (COL1A1, PDGFRB) (in C10 and C13), and cell cycle genes (CCNA2, MKI67) (in C9 and C13) across clusters. **b**, Violin plots showing the expression levels of representative marker genes across all clusters (C0–C14), supporting cluster annotation into EC clusters (C0–C9, C11, 12, and C14), fibroblast/pericyte clusters (C10 and C13), and cell cycle–associated clusters (C9 and C13) are indicated. **c**,**d**, Violin plots showing the expression patterns of ANGPT2 and CLDN5 across experimental conditions (WT_PF, WT_noPF, SWS_PF, and SWS_noPF; see Figure 2B for definitions), indicating the changes specific to the SWS_PF condition. **e**, Over-representation analysis of differentially expressed genes in SWS_PF versus SWS_noPF. Significantly enriched pathways were shown, including the Notch signaling pathway and an apoptosis-related Notch3-associated network, both downregulated in SWS_PF. Dot size indicates overlap percentage, and color represents −log_10_(q-value). **f**, Quantification of cluster composition across experimental conditions, showing the distribution of cells from each condition within each cluster. Cell numbers in each condition were downsampled to enable direct comparison across samples. The proportion of cells in clusters C2, C3, and C7 was decreased in SWS, whereas clusters C9 and C13 were relatively enriched. **g**, Cell proliferation analysis using CFSE labeling. MFI quantification demonstrated increased cell division in SWS-derived cells compared to WT, as indicated by reduced MFI (see right schematic). Statistical significance was determined by an unpaired two-tailed Student’s t-test (n = 3 per group). EC, endothelial cell. CFSE, Carboxyfluorescein succinimidyl ester. MFI, mean fluorescence intensity.

**Extended Data Fig. 3.**
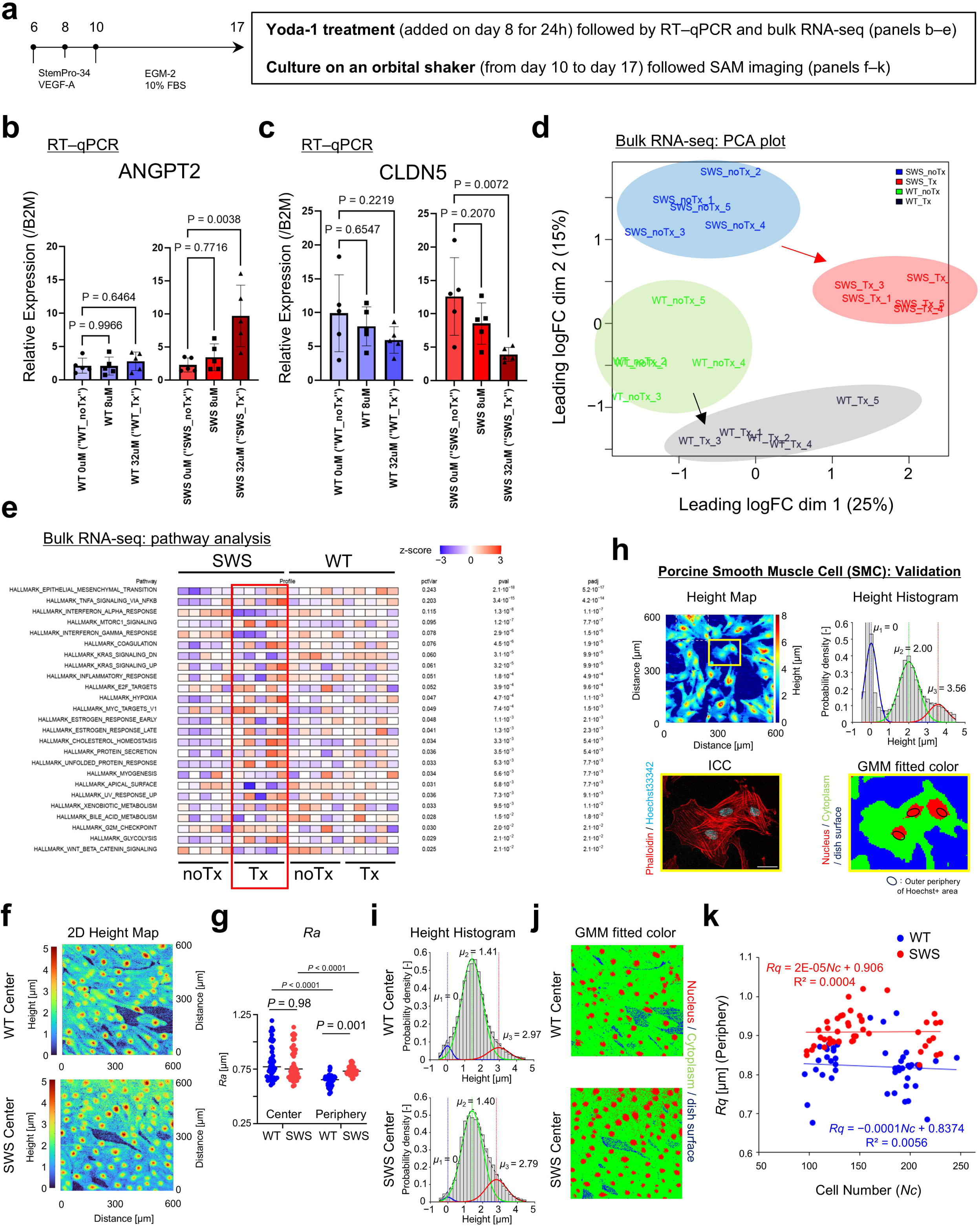
Yoda-1–induced transcriptional responses and SAM-based morphological analyses. **a**, Experimental design of Yoda-1 treatment and orbital shear stress culture. **b**, **c**, RT-qPCR results of ANGPT2 and CLDN5 following Yoda-1 treatment. Statistical significance was determined by one-way ANOVA followed by Dunnett’s multiple comparisons test (n = 5 per group). **d**, PCA of bulk RNA-seq data showing distinct transcriptional responses to Yoda-1 treatment (Tx) (32 µM) across conditions (n = 5 per condition). **e**, Pathway analysis of differentially expressed genes. In SWS ECs, Yoda-1 Tx (32 µM) was associated with activation of inflammatory pathways, including TNFα signaling and mTORC1 signaling, and suppression of apical surface–related pathways. **f**, SAM-based height profiles at the center region of the culture dish, showing minimal differences between WT and SWS ECs. **g**, Quantification of surface roughness (Ra) at the center and peripheral regions of the culture dish. Statistical significance was evaluated by two-way ANOVA followed by Šídák’s multiple comparisons test. Lines indicate the median. **h**, Validation of GMM-based height classification. Representative SAM-derived height map (top left) and corresponding height distribution histogram (top right) from porcine SMCs were shown. The nuclear component identified by GMM fitting (red, bottom right) largely overlaps with Hoechst-positive nuclear regions (blue, bottom left and blue circles in bottom right), supporting the validity of this approach. Scale bar represents 30 µm. **i**, **j**, GMM-based height decomposition in WT and SWS ECs at the center regions. **k**, Correlation analysis between cell number (density) and surface roughness (Rq) at the peripheral regions, showing no significant association in either WT or SWS conditions. SAM, scanning acoustic microscopy. SMC, smooth muscle cell. GMM, gaussian mixture model.

**Extended Data Fig. 4.**
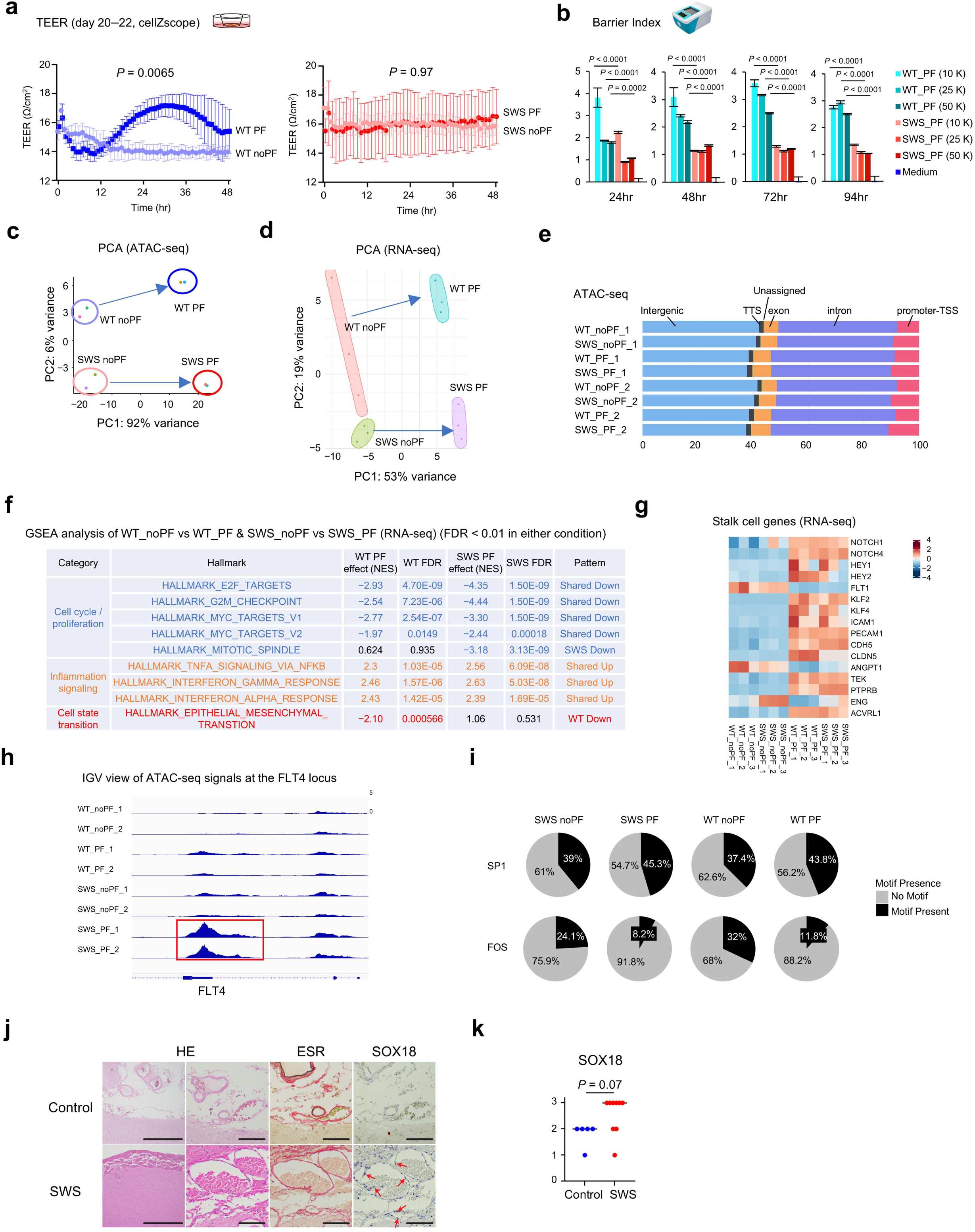
Supporting data for flow-induced functional and multi-omics changes. **a**, TEER measurements using CellZScope. PF enhanced barrier function in WT ECs, whereas SWS ECs showed no discernible PF-dependent change. Data were presented as mean ± SD and were analyzed by two-way repeated measures ANOVA to assess the effects of genotype and time, as well as their interaction. **b**, Barrier index assessed across different seeding densities and time points after PF. Impaired barrier integrity in SWS ECs was consistently observed across all conditions. Data were presented as mean ± SD. Statistical significance was evaluated by two-way ANOVA followed by Šídák’s multiple comparisons test. **c**, **d**, PCA of ATAC-seq (**c**) and RNA-seq (**d**) data showing separation of samples primarily by PF exposure, with more pronounced divergence between WT and SWS under PF conditions. **e**, Genomic distribution of ATAC-seq peaks. SWS ECs under PF showed increased chromatin accessibility in promoter/TSS-associated regions. **f**, GSEA of RNA-seq data comparing PF versus noPF conditions in WT and SWS ECs. Cell cycle–related pathways were consistently downregulated in both groups, whereas inflammatory pathways were upregulated. The EMT pathway shows divergent regulation, being suppressed in WT but not in SWS. NES and FDR values were shown. **g**, Expression of stalk cell–associated genes, showing consistent upregulation in response to PF in both WT and SWS ECs. **h**, Chromatin accessibility tracks of FLT4 showing a PF-induced increase selectively in SWS ECs. **i**, Transcription factor motif accessibility analysis indicating increased prevalence of SP-family motifs and reduced AP-1 motif accessibility in SWS ECs. **j**–**k**, Validation in patient tissue samples. Expression of lymphatic- and tip cell-associated genes was increased in SWS lesions compared to controls. Statistical significance was determined by the Mann-Whitney U test for non-parametric analysis of scoring data. Lines indicate the median. TEER, transendothelial electrical resistance. NES, normalized enrichment score.

**Extended Data Fig. 5.**
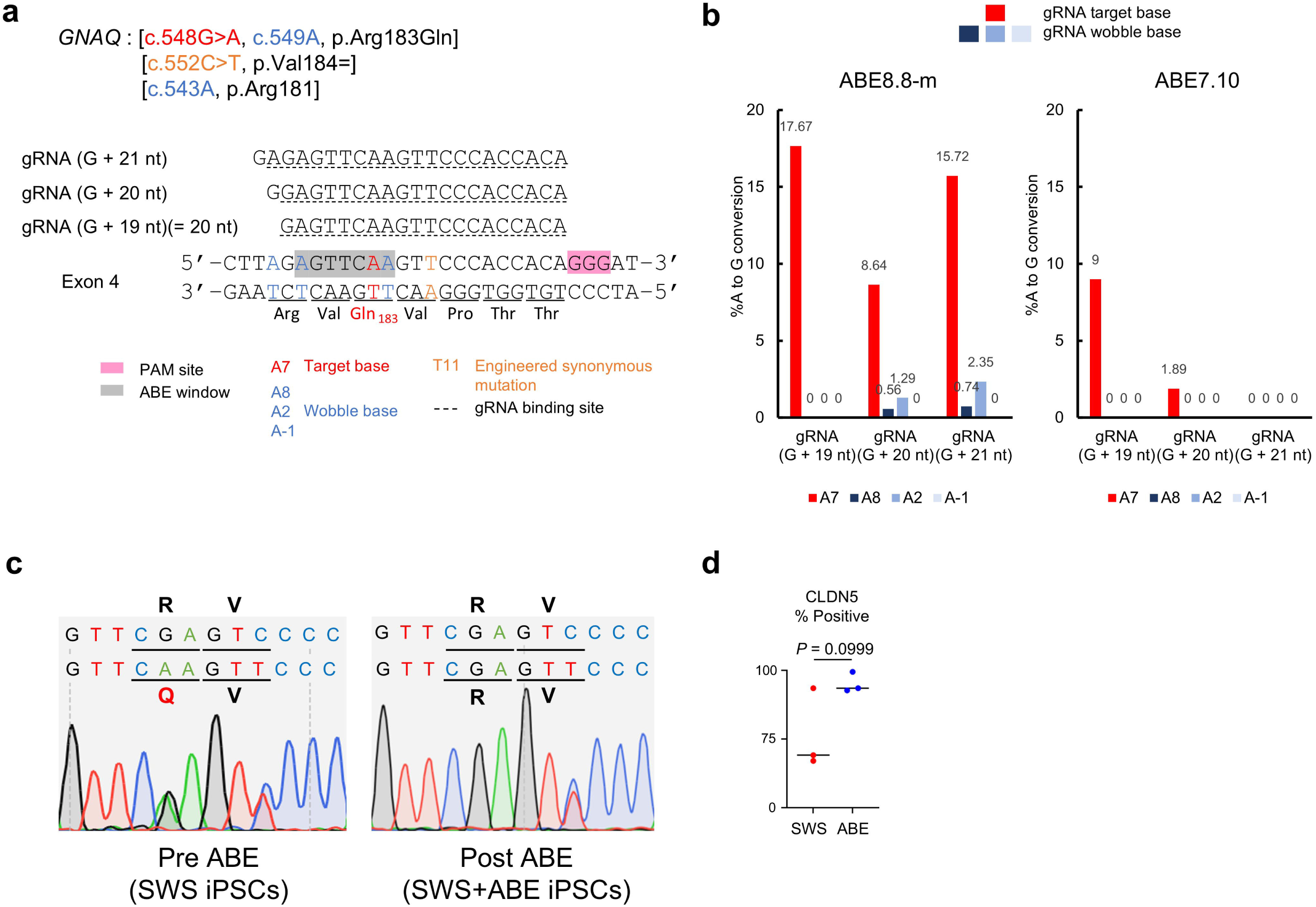
Optimization and validation of adenine base editing for precise correction of the GNAQ mutation. **a**, Design of gRNAs targeting the *GNAQ* locus for adenine base editing. Three gRNAs (G + 19 nt, G + 20 nt, and G + 21 nt) were designed to position the target adenine within the optimal editing window. **b**, Editing efficiencies of A-to-G conversion at target and bystander positions using combinations of gRNAs and ABE variants (ABE7.10 and ABE8.8-m) in SWS iPSCs. Efficient and specific editing was achieved, with minimal bystander activity depending on gRNA design. **c**, Representative sequence analysis of SWS iPSCs before and after ABE correction. The pathogenic c.548G>A mutation was precisely reverted to the wild-type sequence, with no nonsynonymous alterations detected. **d**, Quantification of CLDN5 expression levels in perfused endothelial cells derived from ABE-corrected SWS iPSCs. A trend toward restoration of CLDN5 expression was observed following genome correction. Statistical significance was determined by an unpaired two-tailed Student’s t-test (n = 3 per group). Lines indicate the median. gRNA, guide RNA.

## Methods

### Human induced pluripotent stem cell (iPSC) culture

Human iPSCs were maintained as described previously^1^. Briefly, 409B2 human iPSCs (RIKEN BRC) were cultured on human Lamin-511 (iMatrix-511 silk, Nippi) coated 6-well tissue culture plate (0.5 μg/cm^2^) in StemFit AK02N (AJINOMOTO) medium. For passaging, cells were detached by treatment with TrypLE Select (Thermo Fisher Scientific) at 37 °C for 10 min, followed by gentle mechanical dissociation with a pipette. Typically, 1–2× 10^4^ cells per cm^2^ were seeded on each passage in media containing ROCK inhibitor (Y-27632, Selleck). After 1 day culture, the medium was changed without Y-27632. Five to seven days after plating, the cells reached 80–90% confluency and were again prepared for passage. To make frozen stocks, 2–4× 10^5^ iPSCs were resuspended in STEMCELLBANKER (Takara) and transferred to a cryogenic tube. Stock vials were defrosted onto iMatrix-511 silk coated 6-well plates in StemFit AK02N medium containing Y-27632.

### Design and construction of CRISPR/Cas9 plasmids and donor vector

GNAQ gene- and protospacer sequence of GFP (ps1)-specific gRNA oligos were cloned into pX330-U6-Chimeric_BB-CBh-hSpCas9 (Addgene #42230) (pX-GNAQ and pX-ps1). To derive homology arm left and right (HA-L and HA-R) for GNAQ gene editing, 615 bp and 1544 bp regions were PCR amplified from 409B2 iPSC genomic DNA. Disease-causing mutation (c.548G>A) and synonymous (silent) mutation (c.552C>T) were integrated in the primer of HA-L, and ps1 protospacer were integrated in the primer overhang of HA-L and HA-R. PacI-, ScaI-, ClaI- treated pCAG-mCherry-IRES-puro-pA plasmid (KW154; unpublished) fragment, Acc65I-treated pCAG-eGFP-pA plasmid (KW991; Kim et al., 2018^2^) fragment, HA-L, and HA-R were cloned into one plasmid by employing InFusion Cloning (Clontech). All PCR-amplified regions were verified by sequencing.

### Generation of Sturge-Weber syndrome human iPS cells

GNAQ gene targeting was carried out as described previously^2,3^, with some modifications. Briefly, nuclease expression vector (0.5 µg pX-GNAQ) and the donor vector described above (1.5 µg) were transfected by Neon Transfection System (Thermo Fisher Scientific) system into 2 × 10^5^ cells in single-cell suspension under the condition of one pulse of 1400 V for 20 ms. Electroporated iPSCs were plated at a density of 1.5–5× 10^4^ cells on iMatrix-511 coated 6-well plate in StemFit media containing Y-27632. On day 4, Y-27632 was removed and selection with 0.5 µg/mL puromycin (Sigma-Aldrich) was initiated. On day 14, colonies were isolated manually with a micropipette and clones that underwent homology-directed repair (HDR) with the other allele remaining intact (Extended Data Fig. 1b) were identified by sanger sequencing and expanded further for the next step.

To induce cassette excision, 2 µg of pX-ps1 expression vector was transfected by Neon electroporation into 2 × 10^5^ cells in single-cell suspension, after which cells were plated in 6-well plate in StemFit medium containing Y-27632. On day 8, cassette-excised mCherry negative cells were subjected to fluorescence-activated cell sorting (FACS). Recovered mCherry negative cell populations were later seeded at low density (120–600 cell per 35 mm dish) and single-cell derived colonies were isolated manually. Finally, clones which underwent accurate selection cassette excision were identified by sanger sequencing established as Sturge-Weber syndrome specific iPSCs (“SWS iPSCs”).

### Endothelial cell (EC) differentiation

EC differentiation was performed as described previously^4^. Typically, 30–37× 10^4^ cells per cm^2^ were seeded on growth-factor reduced Matrigel (Corning) in AK02N media containing Y-27632. After 24 h, the medium was replaced with the N2B27 medium (1:1 mixture of DMEM/F12 and Neurobasal medium supplemented with N2 and B27 minus vitamin A; all Thermo Fisher Scientific) with 1µM CP21R7 (Selleck) and 25 ng/ml BMP4 (R&D systems).

After 3 days, the medium was replaced by StemPro-34 SFM (ThermoFisher Scientific) supplemented with 200 ng/ml VEGF (PeproTech) and 2µM forskolin (Sigma), refreshed the following day. On day 6 of differentiation, cells were dissociated with Accutase (Sigma) and enriched for CD34^+^ cells using CD34 MicroBeads (Miltenyi Biotec). The CD34-enriched ECs were either cryopreserved in STEMCELLBANKER (Takara) or plated onto human fibronectin (Sigma)-coated dishes at 50–100× 10^4^ cells per cm^2^ in StepPro-34 SFM supplemented with 50 ng/ml VEGF-A. The medium was changed every other day. On day 9 or 10 of differentiation, ECs were dissociated with Accutase. Cells were then either used for vessel-on-a-chip experiments or replated onto rat tail collagen I (Corning)-coated dishes at 20–25 × 10^4^ cells per cm² in EGM-2 medium (Lonza) supplemented with 10% fetal bovine serum (NICHIREI) for orbital shaker-based culture, depending on the downstream application.

### Microfluidic device

A polydimethylsiloxane (PDMS)-based two-layer, three-channel microfluidic device, fabricated using soft lithography, was used. The channels consisted of three parallel microchannels (Channel 1 and 3 for culture medium, Channel 2 was 1,050 µm in width and separated from the adjacent channels by hexagonal micropillars (100 µm in width, 100 µm in height, and 100 µm spacing between pillars). The micropillar structures were fabricated by a photolithography using SU8 2100 (MicroChem, USA) on a silicon wafer. The features were transferred to the PDMS (Silpot, Dow Corning Toray Co. Ltd.) top layer using soft lithography. Gel injection inlets and medium reservoirs (diameter = 2 mm) were punched into the PDMS layer using biopsy punches (sterile dermal biopsy punch; Kai Industries). The PDMS top layer was then irreversibly bonded to a glass coverslip (Matsunami Glass; 24 x 24 mm2) and baked at 120 °C overnight. The device was sterilized by UV treatment before the experiments.

### Vessel-on-a-chip

A fibrin-collagen gel solution was prepared by combining fibrinogen (Sigma), neutralized rat tail collagen I (Corning), and aprotinin solution (Sigma). Human iPSC-derived ECs or ECFCs (1.6–1.7 ×10^7^ cells/mL) were suspended in EGM2 supplemented with 50 ng/mL VEGF-A. The cell suspension was supplemented with thrombin (1 U/mL; Sigma) to initiate fibrinogen gelation and then mixed with the gel solution at a ratio of 1:1 to achieve final concentrations of 2.5 mg/mL fibrinogen, 0.2 mg/mL collagen, and 0.15 U/mL aprotinin with 8.5 ×10^6^ cells/mL in the mixture. The suspension (20 μL), which exceeded the actual volume of channel 2, was quickly injected into the channel to ensure complete filling. After gel polymerization at 37 °C for 15–30 min, channels 1 and 3 and the wells were filled with EGM2 with 50 ng/mL VEGF-A. Medium was replaced every other day. From day 5 onward, the culture medium was switched to EGM2 without VEGF-A. Perfusion culture was initiated between days 7 and 8, while non-perfusion devices were maintained under static conditions without perfusion.

### Perfusion setup

A ring pump (Aqua Tech) was connected to the ports of channels 1 and 3. To assess lumen formation and vessel connectivity, a perfusion assay was performed by introducing EGM-2 supplemented with 10% FBS and 5 μM rhodamine dextran (70,000 MW; Sigma-Aldrich). The fluorescent tracer was used to confirm the presence of continuous, perfusable vascular structures. For perfusion culture experiments, the pump withdrew medium from channel 3 and delivered it into channel 1 at a flow rate of 1 μL/min, which was expected to produce low-to-intermediate physiological shear stress (approximately 0–10 dyn/cm²), consistent with previously reported microvascular and venous shear stress ranges^5^, based on COMSOL simulations using a comparable microvascular chip system^6^. Perfusion was performed overnight between days 7 and 8 of culture.

### Single-cell RNA-sequencing (scRNA-seq)

Perfused and non-perfused vessel-constituting cells were dissociated using 50 FU/mL (fibrin degradation units) nattokinase (NSK-SD; Japan Bio Science Laboratory Co., Ltd.) in PBS containing 1 mM EDTA (Thermo Fisher Scientific) for 10 min, followed by treatment with Trypsin-EDTA for an additional 10 min. The retrieved cells were resuspended in EGM-2 medium. The cell suspension was then centrifuged at 1,000 rpm for 5 min, and the cell pellet was resuspended in 20–25 μL of EGM-2. Cell viability was assessed using trypan blue staining, and only samples with viability > 80% were used for downstream processing. The scRNA-seq libraries were prepared using the Chromium Next GEM Single Cell 3′ Reagent Kits v3.1 (10x Genomics) according to the manufacturer’s instructions. Briefly, ∼4,000–5,000 cells per sample were loaded onto a Chromium Controller (10x Genomics) for single-cell partitioning and barcoding, followed by cDNA amplification and library construction. Library quality was assessed using an Agilent Bioanalyzer (Agilent Technologies) and Qubit Fluorometer (Thermo Fisher Scientific). Sequencing was performed on a NovaSeq 6000 system (Illumina) to generate paired-end reads, with a sequencing depth of approximately 84,000–159,000 reads per cell. Raw sequencing data were processed using the Cell Ranger pipeline (version 6.1.2, 10x Genomics) with default parameters. Reads were aligned to the human reference genome (GRCh38), and gene expression matrices were generated for downstream analysis.

### Single-cell RNA-seq (scRNA-seq) data processing and analysis

Raw gene expression matrices generated by Cell Ranger were imported into Seurat (version 4.3.0)^7^ in R (version 4.2.2) for downstream analysis. Cells with fewer than 3,000 or more than 8,000 detected genes (nFeature RNA), as well as cells with >15% mitochondrial gene expression, were excluded to remove low-quality or potentially stressed cells. For each dataset, gene expression was normalized and scaled using the default Seurat workflow. Highly variable genes were identified, and principal component analysis (PCA) was performed for dimensionality reduction. Multiple samples were integrated using the Seurat integration workflow to correct for batch effects. The integrated data were used for downstream clustering and visualization. Cells were clustered using a graph-based clustering approach implemented in Seurat. The total number of cells per sample and the number of cells assigned to each cluster are summarized in Supplementary Table 1, together with cluster proportions normalized to the total number of cells in each sample. Uniform Manifold Approximation and Projection (UMAP) was applied for visualization of cell populations. Cell clusters were annotated based on the expression of known marker genes, including endothelial cell markers, fibroblast/pericyte markers, and proliferation markers. Differential gene expression analysis between conditions was performed using the Seurat FindMarkers function. To identify enriched biological pathways, over-representation analysis^8^ was performed on the identified DEGs using the clusterProfiler package, with reference to the KEGG and Reactome databases. The statistical significance of the enrichment was determined using a hypergeometric test followed by Benjamini-Hochberg FDR correction. The accession number for scRNA-seq data reported in this study is GSE330493.

### CFSE proliferation assay

WT and SWS iPSCs were labeled with carboxyfluorescein succinimidyl ester (CFSE; Thermo Fisher Scientific) according to the manufacturer’s instructions. Briefly, cells were incubated with CFSE at a final concentration of 1 μM for 20 min at 37°C, followed by quenching with AK02N medium. After washing, labeled cells were cultured under the indicated conditions. CFSE dilution, reflecting cell proliferation, was analyzed by flow cytometry (SA3800, SONY) at Day 0 and Day 2. Mean fluorescence intensity (MFI) was quantified, and values at Day 2 were normalized to those at Day 0 to calculate relative MFI for comparison between groups.

### Yoda-1 treatment

On day 8 of culture, Yoda-1 (Selleck), a pharmacological agonist of PIEZO channels, was added to StemPro-34 medium at final concentrations of 0, 8, or 32 μM, and cells were incubated for 24 h. Following treatment, cells were dissociated and subjected to downstream analyses. RT–qPCR was performed for samples treated with 0, 8, and 32 μM Yoda-1, whereas bulk RNA-seq was conducted for samples treated with 0 or 32 μM Yoda-1.

### RNA extraction, cDNA synthesis and RT–qPCR

The total RNA from each sample was extracted using RNeasy Plus Micro kit (Qiagen) according to the manufacturer’s instructions. Total RNA (100 ng) was converted to cDNA using SuperScript IV Reverse Transcriptase (Thermo Fisher Scientific). Quantitative PCR (qPCR) was done using PowerUp SYBR Green Master Mix (Thermo Fisher Scientific) and detection was achieved using StepOnePlus Real-Time PCR System (Thermo Fisher Scientific). Primer sequences are listed in Supplementary Table 2. Beta-2-Microglobulin (B2M) was used as a reference gene to calculate transcript abundance of each target gene. The expression level fold-change between sample genes and reference genes was calculated by standard curve, where for every run, a new standard curve was constructed.

### Bulk RNA-seq

The sequence libraries were prepared using a SMART-Seq mRNA or a SMART-Seq v4 Ultra Low Input RNA (Takara Bio) and Nextera XT DNA Library Preparation Kit (Illumina) according to the manufacturer’s protocol.

Sequencing was performed using an Illumina NovaSeq 6000 System with 2 × 150 bp paired-end reads. Reads were mapped to the GRCh38 reference genome using STAR. A GTF file was constructed based on GENCODE 40 annotation. Read counts were generated using featureCounts and analyzed with edgeR. Exploratory data analysis, including Principal Component Analysis (PCA) and generation of heatmaps for global expression patterns and specific gene sets, was conducted using the bulkAnalyseR package. Differentially expressed genes (DEGs) identified via bulkAnalyseR were subsequently subjected to Gene Set Enrichment Analysis (GSEA)^9^ using the clusterProfiler package^10^ in R. Biological pathways and functional categories were enriched based on the Molecular Signatures Database (MSigDB). The accession number for bulk RNA-seq data reported in this study is GSE330654 (Yoda-1) and GSE330603 (orbital shaker).

### Orbital shaker perfusion model

Human iPSC-derived ECs were dissociated on day 10 of differentiation and seeded onto either 35-mm dishes with imaging-compatible membranes or 6-well plates, depending on the downstream application. On day 11, cultures were transferred to an orbital shaker and maintained at 135 rpm in a humidified CO_2_ incubator. Under comparable conditions, computational fluid dynamics simulations have estimated that the peripheral region of the well is exposed to mean wall shear stress values of approximately 0.4–1.2 Pa (4–12 dyn/cm²), within a physiological relevant range, whereas the central region experiences disturbed flow with lower mean wall shear stress values of approximately 0.3–0.8 Pa (3–8 dyn/cm²)^11^.

For imaging-based analyses, ECs were seeded onto 35-mm dishes and cultured under orbital shaking until day 18, followed by immunocytochemistry (ICC), scanning electron microscopy (SEM), and scanning acoustic microscopy (SAM). To prevent medium spillage during orbital shaking, dishes were sealed with a Breathe-Easy membrane (TOHO) that had been sterilized by UV irradiation for 1 h in a clean bench prior to use.

For functional and omics analyses, ECs were seeded onto 6-well plates. Prior to orbital shaking, cells in the central region of the well were mechanically removed, leaving a peripheral cell population to enrich for cells exposed to the shear stress at the periphery. Cultures were then maintained under orbital shaking (135 rpm) from day 11 onward. On day 18, cells were harvested or passaged for downstream analyses, including transendothelial electrical resistance (TEER) measurements (from day 19) as well as RNA-seq and ATAC-sequencing.

### Immunocytochemistry (ICC)

For immunocytochemical analysis, human iPSC-derived ECs and their genome-edited derivative clones were cultured on Lumox dish 35 (Sarstedt) on the orbital shaker as described above. Cells were fixed with 4% paraformaldehyde (PFA), after which the membrane was carefully excised using a scalpel. The membrane samples were subjected to immunostaining with primary antibodies against CD31 (1:100, WM59, BD Biosciences) and Claudin-5 (1:100, ab15106, Abcam), followed by incubation with appropriate secondary antibodies (donkey anti-mouse Alexa Fluor 488 and donkey anti-rabbit Alexa Fluor 546; Thermo Fisher Scientific). Nuclei were counterstained with DAPI. After staining, the membrane samples were mounted onto glass slides using ProLong Gold antifade reagent (Thermo Fisher Scientific). Fluorescence images were acquired using a BZ-X810 microscope (Keyence) or a THUNDER Imager (Leica Microsystems).

### Image Analysis and Quantification

Quantitative analysis of immunofluorescence was performed using ImageJ. Nuclear regions were identified by DAPI staining and defined as ROIs using automated thresholding and watershed segmentation. The mean intensity of Alexa Fluor 546 within each nuclear ROI was measured. Positive cells were quantified based on a cutoff value set at the 5th percentile of the control group’s intensity distribution. Data were collected from independent fields (n = 3–6 per group) and expressed as mean intensity and the percentage of positive cells.

### Scanning electron microscopy (SEM)

PFA-fixed samples cultured on Lumox dish 35 (Sarstedt) were submitted to the Life Science Research Center, Gifu University, Japan, for scanning electron microscopy (SEM) analysis. Sample preparation and imaging were performed according to standard protocols at the facility.

Topography-based evaluation of cell surfaces using scanning acoustic microscopy (SAM) PFA-fixed samples were cultured on the polystyrene film dish (HPS-3805, Honda Electronics) with 50 µm thickness. Quantitative evaluation of cell surface topography was performed using scanning acoustic microscopy (SAM; AMS-50AI, Honda Electronics). The system configuration and acoustic impedance calculation followed our previous work^12^. Measurements were acquired over scanned regions of 600 × 600 µm with a sampling density of 300 × 300 points, encompassing multiple cells per field. PFA fixed cells were measured in PBS, and reflected ultrasonic signals were collected. Time-domain deconvolution was applied using a custom sparse-based algorithm, and the processed signals were spatially reconstructed to generate three-dimensional acoustic impedance maps. Cell height at each point was estimated relative to the substrate interface, yielding spatial height maps of the cell layer. Surface topography was quantified using the root mean square roughness (Rq) and average roughness (Ra). In addition, height distributions were analyzed using a Gaussian mixture model (GMM), and the mean height difference (Δμ) between cell-associated components was used as an index of surface corrugation. Together, these parameters enabled quantitative assessment of multicellular surface heterogeneity. For cross-sectional height analysis, profiles along the direction of flow were extracted from regions containing multiple aligned cells. Three representative positions per region were analyzed, each spanning approximately four cells, to assess spatial variations in cell surface morphology along the flow axis. Detailed analytical procedures are described in the Supplementary Methods.

### cellZScope TEER

ECs for 7 days culture in the orbital shaker (8 x 10^4^ cells) were seeded in cell culture inserts with a pore size of 0.4 μm (Corning) and incubated for 48 h. After changing the media, TEER was measured using a cellZScope (NanoAnalytics) for 48 h.

### Impedance measurement using Maestro Z

Human iPSC-derived ECs cultured under orbital shaking for 7 days were dissociated and seeded onto CytoView-Z plates (Axion BioSystems) at densities of 1 × 10^4^, 2.5 × 10^4^, or 5 × 10^4^ cells per well. Impedance was measured over time using the Maestro Z system (Axion BioSystems) at frequencies of 1 kHz and 41.5 kHz.

Resistance at 1 kHz was used to assess paracellular barrier function (TEER), whereas resistance at 41.5 kHz was used to evaluate transcellular pathways. The barrier index was calculated as ((R_1_kHz + 10) / (R_41.5_kHz + 10)) − 1.

### ATAC-sequencing (ATAC-seq)

Cryopreserved cells were sent to Active Motif to perform the ATAC-seq assay. The cells were then thawed in a 37°C water bath, pelleted, washed with cold PBS, and tagmented as previously described^13^, with some modifications^14^. Briefly, cell pellets were resuspended in lysis buffer, pelleted, and tagmented using the enzyme and buffer provided in the ATAC-Seq Kit (Active Motif cat# 53150). Tagmented DNA was then purified using the MinElute PCR purification kit (Qiagen), amplified with 10 cycles of PCR, and purified using Agencourt AMPure SPRI beads (Beckman Coulter). Resulting material was quantified using the iSeq (Illumina), and sequenced with PE50 on the Illumina sequencing platform.

Reads were aligned using the BWA algorithm (mem mode; default settings). Duplicate reads were removed, only reads mapping as matched pairs and only uniquely mapped reads (mapping quality >= 1) were used for further analysis. Alignments were extended in silico at their 3’-ends to a length of 200 bp and assigned to 32-nt bins along the genome. The resulting histograms (genomic “signal maps”) were stored in bigWig files. Peaks were identified using the MACS 2.1.0 algorithm at a cutoff of p-value 1e-7, without control file, and with the –nomodel option. Signal maps and peak locations were used as input data to Active Motif’s proprietary analysis program, which creates Excel tables containing detailed information on sample comparison, peak metrics, peak locations, and gene annotations. The accession number for ATAC-seq data reported in this study is GSE330603.

### TOBIAS analysis

TOBIAS was used to perform ATAC-seq footprinting of known transcription factors^15^. In brief, TOBIAS corrects for Tn5 bias, calculates footprint scores from the distribution of Tn5 insertions and compares these with all predicted binding sites based on position weight matrices. All TOBIAS analyses were run with default parameters and the HOCOMOCO version 11 database as reference (https://doi.org/10.1093%2Fnar%2Fgkx1106).

### Patient sample and immunostaining

For histological analysis, tissue samples were fixed in 4% PFA overnight at 4°C and subsequently stored in 70% ethanol at 4°C. Hematoxylin and eosin (HE) staining and immunohistochemistry (IHC) were performed on 3-μm-thick paraffin-embedded sections. Sections were deparaffinized in xylene and rehydrated through graded ethanol. Endogenous peroxidase activity was quenched with 0.3% hydrogen peroxide in methanol for 20 minutes. Antigen retrieval was performed in a pressure chamber using Tris–EDTA buffer (7.4 mM Tris-HCl, 1 mM EDTA-2Na, pH 9.0). Primary antibodies were as follows: AQP1 (20333-1, Proteintech, 1:100), NR2F2 (EPR18443, Abcam, 1:150), and SOX18 (sc-166025, Santa Cruz Biotechnology, 1:50).

Secondary detection was performed using the Histofine Simple Stain System (Nichirei Biosciences) for 1 h. Peroxidase activity was visualized with DAB–H_2_O_2_. Images were acquired using a Keyence BZ-X700 microscope and processed with ImageJ software. For Elastica Picrosirius Red (ESR) staining, sections were deparaffinized, rehydrated, and treated with 1% hydrochloric acid in 70% ethanol. Sections were stained with Weigert’s resorcin-fuchsin for 40–50 minutes to visualize elastic fibers, followed by counterstaining with Weigert’s iron hematoxylin for 3–5 minutes. Sections were then stained with Picrosirius Red solution for 15 minutes to visualize collagen fibers. In ESR staining, collagen fibers appear red, elastic fibers black, and muscle fibers yellow. All reagents for special staining were purchased from Muto Pure Chemicals Co., Tokyo, Japan. For semi-quantitative analysis of protein expression, immunostaining intensity was scored by the observer blinded to the experimental groups. The expression levels were categorized into a four-point scale: 0 (none), 1 (weak), 2 (moderate), and 3 (strong).

### Plicamycin treatment

For pharmacological inhibition of SP1/2 transcription factors, plicamycin (Selleck) was applied to ECs cultured under orbital shaker conditions as described above. Cells were assigned to three experimental conditions: (i) disease model (0 nM), (ii) prevention model (2 nM from the start of orbital shaking), and (iii) treatment model (0 nM for the first 5 days followed by 2 nM for the final 2 days). After 7 days of orbital shaker culture, cells were collected, and CLDN5 expression was evaluated by RT–qPCR.

### Adenine base editing

To optimize base editing conditions, sgRNAs of varying lengths were designed to match the editing window, and two adenine base editors (ABE7.10 and ABE8.8-m) were evaluated. SWS iPSCs were co-transfected with sgRNA and ABE constructs using the Neon Transfection System (1400 V, 20 ms, 1 pulse). After transfection, cells were subjected to puromycin selection for 24–72 h, followed by genomic DNA extraction. Target regions were amplified by PCR and subjected to amplicon sequencing. FASTQ files were analyzed using CRISPResso2 (https://crispresso2.pinellolab.org/)^16^. Using the optimized conditions, SWS iPSCs were transfected as described above, and single-cell-derived clones were established following antibiotic selection. Genomic DNA from each clone was PCR-amplified, and restriction fragment length polymorphism analysis using TaqIv2 was performed to screen for corrected alleles based on digestion patterns. Selected clones were differentiated into endothelial cells and cultured under orbital shaker conditions as described above. CLDN5 expression was then evaluated by immunocytochemistry.

### Study approval

The study protocol was approved by the Institutional Review Board (IRB) of Nara Medical University (approval number: G151), which served as the central ethics committee for this multi-institutional collaboration. Human samples including umbilical cord blood (Nara Medical University), surgical specimens from SWS patients (National Center of Neurology and Psychiatry and Niigata University), and post-mortem tissues (Mie University) were obtained following written informed consent from all the participants or their legal guardians, in accordance with the Declaration of Helsinki. The human iPS cell line 409B2 (RIKEN BRC) and its genome-edited variants were also utilized under the same ethical approval (G151) and relevant transfer agreements.

### Statistical Analysis

Statistical analyses were performed using GraphPad Prism 9 (GraphPad Software). For comparisons between two groups, an unpaired two-tailed Student’s t-test was used for normally distributed data, while the Mann-Whitney U test was applied for the analysis of immunostaining scores. For multiple group comparisons, one-way analysis of variance (ANOVA) followed by Dunnett’s multiple comparisons test or two-way ANOVA (or repeated measures two-way ANOVA where appropriate) followed by Šídák’s or Tukey’s post hoc tests were employed, as indicated in the figure legends. Specifically, cell physiological assays (Maestro and cellZscope) were analyzed using two-way repeated measures ANOVA to account for temporal or matched-sample dependencies. Data are presented as mean ± SD from at least three independent experiments. P-value < 0.05 was considered statistically significant.

