## Supplemental Method for "Dysregulated flow responses drive pathogenesis in a cellular model of Sturge–Weber syndrome"

**Supplementary Methods**

**Three-dimensional reconstruction and topography-based analysis using scanning acoustic microscopy**

**S1. Scanning acoustic microscopy measurement and signal processing**

Scanning acoustic microscopy (SAM) was used to evaluate the surface topography of cultured cells. The overall experimental system configuration was similar to that described in our previous report^1^. Measurements were performed using a commercial scanning acoustic microscopy system (AMS-50AI, Honda Electronics) equipped with a 280-MHz transducer (HTD400-09, Honda Electronics). Pure water was used as the coupling medium. Reflected ultrasonic signals were acquired from cells adhered to the bottom of a dedicated culture dish designed for acoustic measurements (HPS-3805, Honda Electronics).

Cells were chemically fixed with 4% paraformaldehyde (PFA) and maintained in phosphate-buffered saline (PBS) during measurements. For each acquisition, a scanning area of 600 × 600 µm was sampled at 300 × 300 measurement points, covering multiple cells within each scanned region. Accordingly, the analysis focused on surface topography at the multicellular level rather than on individual cell morphology.

The acquired signals were processed using time-domain deconvolution to resolve reflections along the depth direction. This analysis employed a sparse-based approach, based on the physical assumption that acoustically reflecting interfaces are discretely distributed along the depth axis. As a result, depth-dependent acoustic impedance profiles were obtained at each scanning position.

Label-free, non-destructive three-dimensional imaging approaches based on acoustic signals have been reported previously^2,3^. The analysis employed here was developed independently to address the characteristics of ultrasonic reflection signals and the experimental configuration used in the present study.

**S2. Three-dimensional reconstruction and definition of cell height**

Depth-dependent acoustic impedance profiles obtained at individual scanning points were spatially arranged according to their corresponding lateral scan positions, enabling reconstruction of three-dimensional acoustic impedance distributions.

Within the reconstructed three-dimensional distributions, the substrate–medium interface and the medium–cell interface were identified. Cell height at each measurement point $(x,y)$, denoted as $H(x,y)$, was defined as the distance between these two interfaces:

$$H(x,y)=z_{\text{cell}}(x,y)-z_{\text{sub}}(x,y)$$

(1)

where $z_{\text{sub}}$ represents the depth of the substrate–medium interface and $z_{\text{cell}}$ represents the depth of the medium–cell interface. Based on this definition, height maps $H(x,y)$ were constructed for each scanned region. These height maps served as the basis for evaluating surface topography across scanned regions containing multiple cells.

**S3. Quantitative evaluation of surface topography**

To quantify overall surface topography within each scanned region, the root mean square roughness (*Rq*) was calculated from the height map as follows:

$$Rq=\sqrt{\frac{1}{N}\sum_{i=1}^{N} \left( H_{i} - \bar{H} \right)^{2}}$$

(2)

where $H_{i}$ is the height at the $i$-th measurement point, $\bar{H}$ is the mean height within the scanned region, and $N$ is the total number of measurement points.

To further characterize the structural nature of surface corrugation, the height distribution within each scanned region was statistically decomposed using a three-component Gaussian mixture model (GMM; *k* = 1–3). The three components corresponded to the substrate–medium interface (*k* = 1), a lower cell-associated height component (*k* = 2), and a higher cell-associated height component (*k* = 3). Here, $\mu_{\text{1}}$ represents the height of the substrate–medium interface and was fixed at 0 in this study. $\mu_{\text{2}}$ and $\mu_{\text{3}}$ represent the mean heights of the lower and higher cell-associated Gaussian components, corresponding to basal cytoplasmic regions and nuclear-associated elevated regions, respectively. The boundary between basal cytoplasmic regions and nucleus-associated elevated regions was determined using a GMM-derived height threshold *T*_23_ defined as:

$$T_{23}=\frac{\pi_{3}\mu_{2}+\pi_{2}\mu_{3}}{\pi_{2}+\pi_{3}}$$

(3)

where $\mu_{k}$ and $\pi_{k}$ denote the mean height and mixing proportion of component *k*, respectively. This formulation corresponds to an inverse-mixing-ratio interpolation between the two components, such that the component with the larger proportion occupies a wider spatial region, enabling deterministic segmentation even when distributions overlap.

The mean values of these components were used as representative heights. The difference between the mean heights of the higher and lower cell-associated components was defined as Δ*μ*:

$$\Delta\mu=\mu_{\text{3}}-\mu_{\text{2}}$$

(3)

By combining *Rq*, *Ra*, and Δ*μ*, both the magnitude and structural characteristics of surface topography within multicellular regions were quantitatively evaluated.

1	Ujihara, Y., Watanabe, S., Morodomi, S., Sugita, S. & Nakamura, M. Probing mechanical properties through acoustic impedance in cultured smooth muscle cells with cytoskeletal disruption using scanning acoustic microscopy. *Journal of Biorheology* **38**, 112-119 (2024). <https://doi.org:10.17106/jbr.38.112>

2	Bagus Prastika, E. *et al.* Time and frequency domain deconvolution for cross-sectional cultured cell observation using an acoustic impedance microscope. *Ultrasonics* **119**, 106601 (2022). <https://doi.org:10.1016/j.ultras.2021.106601>

3	Hozumi, N., Yoshida, S. & Kobayashi, K. Three-dimensional acoustic impedance mapping of cultured biological cells. *Ultrasonics* **99**, 105966 (2019). <https://doi.org:10.1016/j.ultras.2019.105966>
